# Development of the Follow-Up Discourse Observation Protocol (FUDOP) for characterizing instructor active-learning follow-up behaviors

**DOI:** 10.1101/2024.08.20.608824

**Authors:** Xinjian Cen, Maci Kight, Rachel Lee, Petra Kranzfelder, Stanley M. Lo, Jeffrey Maloy, Melinda T. Owens

## Abstract

Instructors can provide feedback to their class in multiple ways. One way is through their follow-up behaviors, which are the specific strategies instructors implement after active learning activities. These behaviors could play an important role in student learning as students can receive feedback from the instructor. However, there is little research on the effects of different types of follow-up behaviors. Follow-up after active learning can be seen as a form of discourse between the instructor and the entire class. Previous researchers developed the Classroom Discourse Observation Protocol (CDOP) to analyze discourse between the instructor and individual students or small groups. We used CDOP as a starting point to develop and validate a new protocol, the Follow-Up Discourse Observation Protocol (FUDOP), to characterize instructional follow-up behaviors to the entire class after active-learning activities. We then used FUDOP to characterize follow-up behaviors of multiple instructors in introductory biology courses at three different universities. We measured consistent differences in these behaviors across instructors but not within instructors, demonstrating that instructors may engage in consistent follow-up behaviors. FUDOP could allow instructors and researchers to better measure and analyze follow-up behaviors and their effects, which could in turn provide guidance to instructors and faculty developers.

## Introduction

In any classroom, many instructors can incorporate active learning, a type of student-centered pedagogy that involves student engagement and interaction through class activities (Driessen et al., 2020). This engagement may take many forms, e.g. individual thinking, writing, or peer discussion, where students collaborate and explain their understanding of concepts to exchange ideas (Auerbach & Schussler, 2017). Active learning facilitates engagement with the material by having students answer questions about the content and interact with other students (Chin, 2006). When compared to traditional lectures, active learning has been shown to increase student performance and decrease failure rates (Dewsbury et al., 2022; Freeman et al., 2014; Theobald et al., 2020). While there are many activities that potentially fall under the umbrella of active learning, some common active learning activities include instances where the instructor prompts students to engage in discussions with their peers (hereafter referred to as peer discussion) and questions that students respond to using classroom response systems (hereafter referred to as clicker questions) as these often allow for instructor feedback following the activity (Freeman et al., 2014; Mazur, 1997).

### Active learning and discourse

Instructors can use teacher discourse moves, an interactive element in a science classroom to increase discourse, as a linguistic resource that can help them increase student engagement and facilitate learning (Bae et al., 2021). Discourse can be a dialogic process: both the instructor and the student are engaged in it, and words are not looked at by themselves but rather in conjunction with their context to decipher their meaning (Mortimer & Scott, 2003). Thus, when students engage in discourse with each other or with the instructor, it is an active process that has the potential to allow students to use their collective knowledge to develop an understanding of the material being taught (Blanton, 2002). The interaction between students and instructors in the classroom can play an important role in students’ learning process, which can affect their understanding and application of the material (Bae et al., 2021). There has been a growing body of research categorizing different kinds of discourse in the college STEM classroom and how they could support students’ learning in different respects (Alkhouri et al., 2021; Bae et al., 2021; Boud & Molloy, 2013; Chin, 2006; Van Booven, 2015; Wood et al., 2018).

Some researchers have broadened their conceptions of discourse to include the discourse students engage in with each other during active learning (Wood et al., 2018). That means that when characterizing the instructor’s approach, one not only needs to consider who speaks when but also the “ideological stance” of the discourse (Ford & Wargo, 2012; O’Connor & Michaels, 2007). One study of discourse in an undergraduate physics course found that even when the instructor’s discourse moves were structurally authoritative, in that the instructor was not looking for direct student replies, they could be ideologically dialogic because they were prompting the students to talk with each other, thus having them engage in dialogue and construct higher-level thinking (Wood et al., 2018).

Most studies of discourse have been done in small K-12 classrooms, where many students have the opportunity to interact with their instructor as part of a whole-class discussion, but these discourse patterns can also be seen in large college classes (Bae et al., 2021). However, the nature of discourse patterns in large college classes can be more complicated because the size of the classroom means there are more types of possible interactions (Wood et al., 2018). That complexity increases if the instructor uses active learning techniques to engage the class in various ways. In these large classrooms, there may be little direct interaction between instructors and individual students, which reduces the opportunities for individualized teacher discourse moves (Blanton, 2002; Chin, 2006). Instead, to interact with students, instructors can use active learning activities, which they may then follow up in various ways to either solicit more student input or to communicate information to them (Mesa & Chang, 2010). In this paper, we claim that instructors’ follow-up to active learning activities can also be considered a form of classroom discourse. Here, we define “follow-up” as any action the instructor takes after an active learning activity that is in response to that activity, as shown in Figure 1. (Note that our definition is broader than the one given in the Classroom Observation Protocol for Undergraduate STEM (COPUS), in which follow-up is solely defined as the content-based feedback on a clicker question or activity given to the entire class (Smith et al., 2013)). We created and validated a protocol, the Follow-Up Discourse Observation Protocol (FUDOP), to characterize these follow-up activities, and used it to describe instructor follow-up patterns in active learning classrooms at several universities.

**Figure 1:**
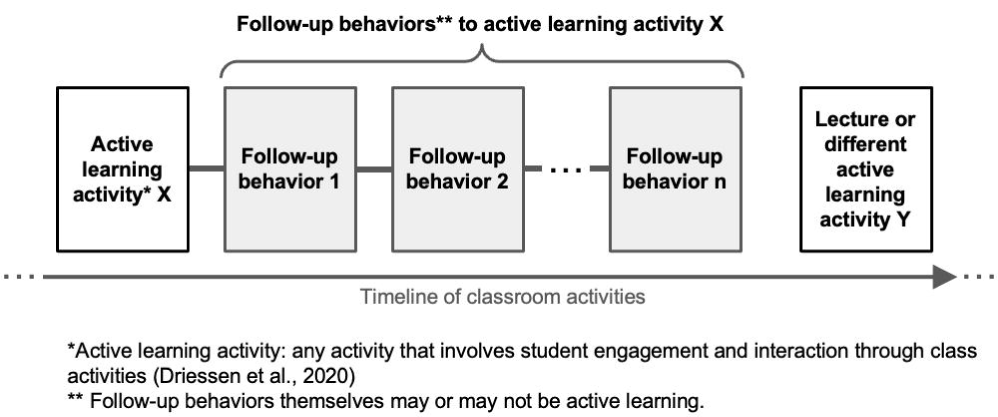
Follow-up in the context of a classroom timeline. Follow-up occurs after an active learning activity, which can happen any time during a class session.

In active learning, instructors often have the opportunity to engage in conversations with individual students or small groups during the active learning activities, which is another opportunity for discourse (Lombardi et al., 2021). Some researchers have analyzed this type of discourse between the instructor and individual students or small groups to create the Classroom Discourse Observation Protocol (CDOP) (Kranzfelder et al., 2019). CDOP quantifies what instructors do and allows for the categorization of teacher discourse moves into 17 codes, which are classified using the “approach” part of Mortimer and Scott’s framework (Alkhouri et al., 2021; Mortimer & Scott, 2003). Five of these codes are “teacher-centric” because they involve the instructor as the dominant voice discussing content, while ten are “student-centric” because the students talk about content (Kranzfelder et al., 2019). Further work with CDOP has revealed that teaching and discourse practices in STEM classrooms vary significantly across discipline and instructor positions but less so based on levels of experience and class sizes (Alkhouri et al., 2021). However, even though the active learning classroom provides more opportunities than a traditional lecture for the instructor to interact with individual students, the large size and layout of many undergraduate classrooms mean that most students will not have the opportunity to have a one-on-one interaction with the instructor during class time.

### Framework: classroom discourse and its importance

Frameworks for characterizing discourse grew out of sociocultural theory, which emphasizes that students make sense of the world through interaction with instructors and peers (Lemke, 2001). Early researchers began to categorize discourse methods used by instructors by classifying them along axes (Mortimer & Scott, 2003). This arises from the fact that different instructors at different times have different reasons for asking questions or initiating communication, which adds meaning to the discourse. The axes of classification can be described by two dimensions: interactive/non-interactive and dialogic/authoritative, resulting in four individual categories. An interactive classroom includes the instructor and students discussing a topic either from one perspective, most notably the instructor’s, or from multiple perspectives if there is an opportunity for students to give their opinion. This method of teaching involves a conversation and a series of questions and answers between students or student and instructor. In contrast, in a non-interactive classroom, only the instructor speaks in more of a lecturing style.

This method of teaching closes off the conversation because the students are not able to engage in conversation with the instructor or reply to other students. A classroom with a dialogic approach takes multiple perspectives into account, including those of the students. In contrast, an authoritative classroom only allows discussion from one perspective. This can be the instructor lecturing from their perspective or asking the class or an individual student to answer a question with a desired response.

The two dimensions are independent. In other words, classrooms can be interactive dialogic, interactive authoritative, non-interactive dialogic, or non-interactive authoritative. For example, if an instructor asks, “What is the first stage of mitosis?” and the class answers, “Prophase,” it is interactive because the students are responding to the instructor’s question, but it is authoritative because the instructor clearly has one “correct” answer in mind. On the other hand, if an instructor includes summaries of the thoughts or ideas students had on a topic as part of a lecture, it is non-interactive because only the instructor is speaking, but it is dialogic because the instructor is including the student’s perspectives. Table 1 shows examples of actions from previous literature that would fit into these four categories (Mortimer & Scott, 2003).

**Table 1:**
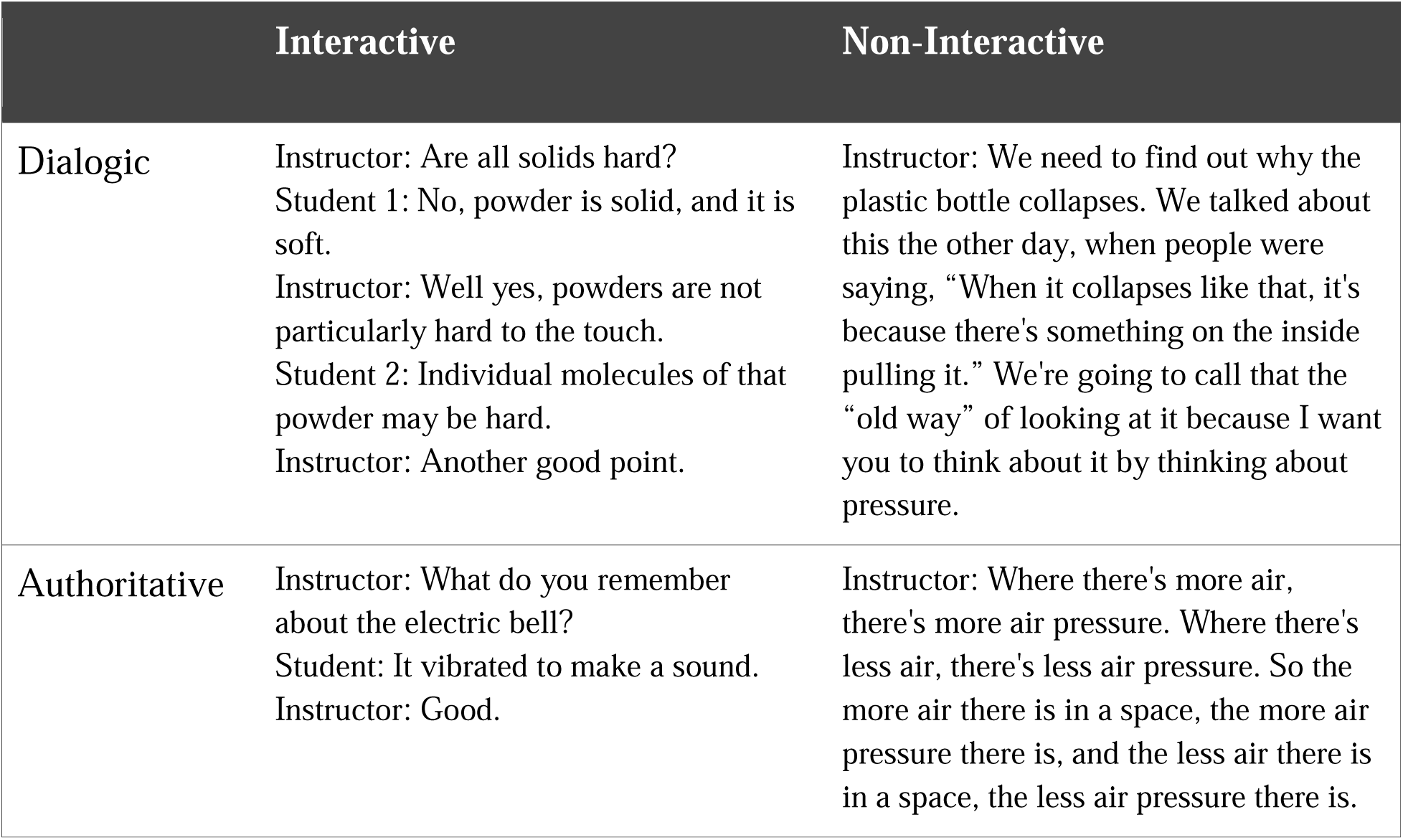
Examples of Discourse Approaches along the Interactive/Non-Interactive and Authoritative/Dialogic Axes. Summarized from Mortimer and Scott, 2003.

|  | Interactive | Non-Interactive |
| --- | --- | --- |
| Dialogic | Instructor: Are all solids hard?<br>Student 1: No, powder is solid, and it is soft.<br>Instructor: Well yes, powders are not particularly hard to the touch.<br>Student 2: Individual molecules of that powder may be hard.<br>Instructor: Another good point. | Instructor: We need to find out why the plastic bottle collapses. We talked about this the other day, when people were saying, “When it collapses like that, it's because there's something on the inside pulling it.” We're going to call that the “old way” of looking at it because I want you to think about it by thinking about pressure. |
| Authoritative | Instructor: What do you remember about the electric bell?<br>Student: It vibrated to make a sound.<br>Instructor: Good. | Instructor: Where there's more air, there's more air pressure. Where there's less air, there's less air pressure. So the more air there is in a space, the more air pressure there is, and the less air there is in a space, the less air pressure there is. |

Patterns of classroom discourse can affect student engagement and learning. The quality of discourse from the instructor directly affects the degree of student learning with student learning increasing as instructors employ discourse moves that encourage engagement (Zhang, 2009). A study of the reasons instructors give for why they use dialogic discourse shows that instructors believe it can enhance cognitive engagement, initiate learning activities, offer relevant feedback, harness student ideas, and prompt students to establish connections (Donham & Andrews, 2023). A detailed analysis of undergraduate math courses found that different discourse practices can indeed either provide or hold back opportunities for students to engage in the classroom. This analysis looked at two different undergraduate instructors teaching two mathematics courses and found that the classroom with an instructor that used more linguistic dialogue led to higher student engagement with the instructor and between each other (Mesa & Chang, 2010). Other papers have found that different discourse patterns do prompt different levels of student thinking. Using IRF instead of IRE, for example, can provide the discursive space encouraging students to engage in a higher level understanding of the concepts, promoting further student input, and deeper thinking beyond simple recall (Bae et al., 2021; Chin, 2006; Van Booven, 2015).

### Follow-up after active learning as a form of discourse

One way that students communicate with instructors in large classes is through follow-up to active learning activities. Instructors have a lot of choices for how to conduct follow-up activities. For example, three instructors may ask a clicker question, and the students all individually select their response. One instructor may choose to call on individual students to explain their thinking, another instructor may tell the students to discuss their reasoning with their neighbors, and the third instructor may simply acknowledge the work the students did during the active learning segment and move on to the next topic without further interaction with the students about the original question (Chin, 2006). In each case, the instructor is engaging in a different type of interaction with the entire class, which gives different types of feedback to the class and allows different students to communicate with the instructor. In other words, the follow-up to the active learning causes there to be discourse between the instructor and the entire class, and the three different instructors are participating in different discourse patterns (Mortimer & Scott, 2003).

Instructor follow-up after active learning has not been extensively studied. In theory, instructor follow-up to active learning activities completes a “feedback loop” that begins with the instructor’s clicker question or other active learning activity. A single active learning activity can even lead to multiple rounds of follow-up, for example, if a student asks a question during follow-up, which the instructor responds to by delivering another question, creating a positive feedback loop. These interactions create an ideologically dialogic classroom that can stimulate student thought (Boud & Molloy, 2013; Wood et al., 2018). Also, instructor follow-up after an active learning activity is an example of instructor instructional feedback, which is a powerful element that influences student learning by delivering focused content exactly when the student is most interested in it (Hattie & Timperley, 2007). In practice, however, there has been less research on how instructors implement follow up in practice and the effects of follow-up. There is literature attesting that different instructors implement follow-up differently. For example, studies on “fidelity of implementation” suggest that instructions following the “peer instruction” method of active learning, where instructors are supposed to use peer discussion to follow up after an individual clicker activity, the instructors often skip the peer discussion (Smith et al., 2011). This individual variation in follow-up preferences may matter for learning. One study found more student learning when an instructor gives an explanation after an active learning activity, compared to when an instructor merely administers the active learning activity and does not follow up with an explanation (Smith et al., 2011). Additional studies using classroom observation protocols have found that within an individual instructor, the instructional behaviors they employ may vary between class sessions, introducing an additional potential source for variation (Stains et al., 2018). However, there is still much more to learn about what instructors do during follow-up and what effects their choices might have on student learning.

In our study, we seek to understand this relatively understudied discourse that occurs between instructors and students following the instance of an active learning activity. We used CDOP as a starting point to develop and validate a new observation protocol, the Follow-Up Discourse Observation Protocol (FUDOP), to characterize instructional follow-up after active learning activities. We wanted to see the extent to which follow-up discourse patterns might be an interesting axis of variation and that might modulate the effect of active learning. This led us to analyze the extent to which these discourse behaviors vary between different instructors by applying FUDOP to characterize the follow-up behaviors of instructors at three different universities. Our specific research questions were:

1. To what extent can we adapt CDOP to categorize discourse behaviors of instructors when following up on active learning activities?
2. Do these discourse behaviors vary between different instructors?
3. Do individual instructors use consistent discourse behaviors across different class sessions?

We hypothesized that this newly developed tool would reveal discourse patterns that might differ significantly between instructors and that for each individual instructor these discourse patterns might vary significantly between class sessions.

## Methods

The study was approved by the Institutional Review Board (IRB) offices of Northwestern University (protocol #STU00042205), the University of California, Los Angeles (protocol #19-001554), and the University of California, Merced (protocol #UCM 2020-3).

### Positionality Statement

Our research benefits from a diverse range of perspectives, reflecting viewpoints of individuals who are on the inside and outside of the educational contexts in our study (Boveda & Annamma, 2023; Holmes, 2020). Author J.M. offers valuable firsthand knowledge as one of the instructors included in our dataset, providing insights into instructional practices and challenges. Additionally, authors M.T.O., S.L., and P.K. bring expertise as biology education researchers actively engaged in teaching large STEM classes similar to those examined in our study, offering valuable insights into broader educational trends and practices. As regular instructors of large undergraduate active learning courses, each of these authors also contributes valuable perspectives regarding instructor motivations for engaging in various discourse moves. As education researchers, they provide perspectives on how FUDOP codes will be understood and used by education researchers in different settings. Author X.C. contributes a student perspective, having experienced some of the courses studied here firsthand. This allows her to contribute context about the student experience during active learning segments and about alignment between specific assessment questions and course learning objectives. Authors R.L. and M.K. provide further student perspectives, having experienced diverse teaching styles within the educational settings similar to our study. Together, these varied viewpoints enrich our research, fostering a comprehensive understanding of the complexities inherent in STEM education.

### FUDOP Development Strategy

First, we designed a protocol that would allow observers to reliably characterize common instructor follow-up behaviors. We created our protocol by adapting codes from the existing, validated CDOP protocol (Kranzfelder et al., 2019). Using CDOP, we began viewing a subset of classroom video recordings, and then iteratively added new codes if follow-up behaviors occurred that were not described adequately by the CDOP-derived codes. This process continued until no new codes were needed to describe classroom practices observed in our sample dataset. After developing and validating the FUDOP protocol, we asked whether we could use it to measure consistent differences in these behaviors between instructors, using a set of instructors from multiple institutions (Figure 2). In the next section, we will expand on our study context and development process.

**Figure 2:**
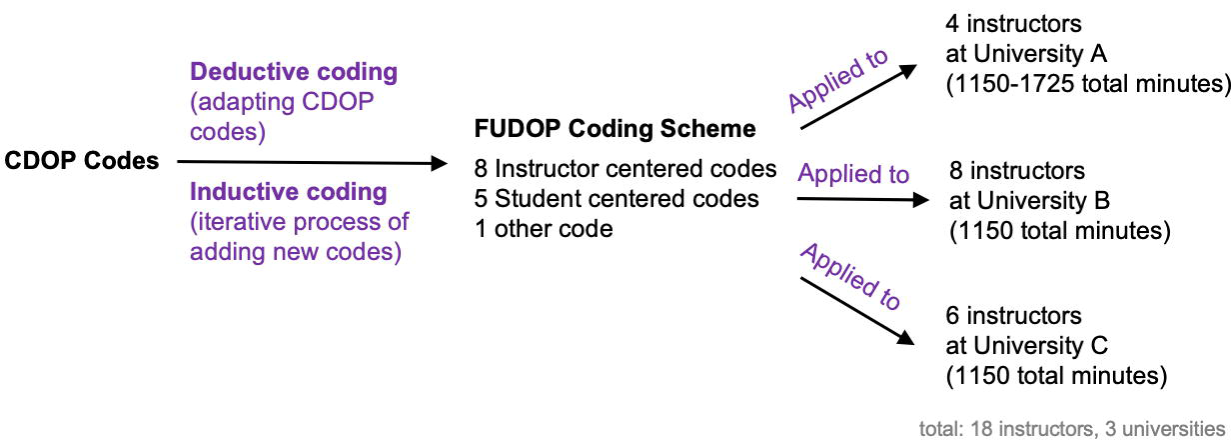
Development of FUDOP.

### Study Context

To develop FUDOP, we obtained a large data set of classroom recordings consisting of 46 individual 50-75-minute in-person lectures taught by 18 different instructors from introductory biology courses at three four-year universities in the United States (Supplemental Table 1). The universities included in this study are diverse in size, institution type, student population, and geography, as described further below and in Supplemental Table 1. We chose to have all the recordings come from classes teaching introductory biology material so that if we found differences between instructors in follow-up behavior, those differences would not be attributable to the content taught or varying disciplinary norms. In all cases, the recordings analyzed came from various points during the term and covered a variety of introductory biology topics.

The introductory biology course analyzed at university A, a public West Coast R1 university, had three offerings of 340-380 students each, taught by four instructors total. From these offerings, we analyzed 23 50–75-minute-long lectures. All instructors were full-time non-tenure-track instructors with multiple years of experience teaching and coordinating the course who have received professional development on implementing active learning. All offerings were taught in the same quarter, and the lectures from university A had the four instructors sharing the same set of slides, assessments, and active learning activities. Thus, any variation in follow-up behavior could be attributed solely to instructor discretion.

The four introductory biology courses analyzed at university B, a private Midwestern R1 university, had eight offerings of 200-250 students, which were taught by different instructors with 0-20+ years of teaching experience. From these offerings, we analyzed 23 50-minute-long lectures. Courses were taught by tenure-track research-focused instructors who were all part of a professional and curriculum development program aiming to develop a learner-focused first-year biology curriculum. Beyond the collaborative program activities, each instructor had autonomy in how they developed and implemented their course curriculum.

The two introductory biology courses analyzed at university C, a public West Coast R2 university, had four offerings of 39-236 students each, taught by six instructors total. University C is also a Hispanic-serving institution (HSI). We analyzed 22 50–75-minute-long lectures. Courses were co-taught by tenure-track research-focused instructors, tenure-track teaching-focused instructors, and non-tenure track instructors with multiple years of teaching experience. All of the instructors developed their lecture slides and in-class activities independently, but coordinated the rest of the course together, including assessments. The details of the number of instructors, courses, course offerings, and lectures analyzed for all universities is summarized in Supplemental Table 1.

### Identifying instances of active learning and follow-up

Consistent with well-established definitions of active learning, we identified instances during class sessions where students engaged and interacted with each other and the instructor through individual thinking, writing, or peer discussion (Driessen et al., 2020). In all courses examined here, the active learning activities consisted of clicker questions, peer discussion, and worksheet questions (the instructor provided written exercises or problems for students to complete individually or in groups). After each active learning activity was presented, a certain amount of time was allotted for students to complete the activity.

We defined instructor follow-up as beginning when the initial activity was finished and the instructor regrouped the class. A single active learning activity often prompted multiple instructor follow-up behaviors. For example, sometimes instructors initiated additional active learning activities like a clicker question or peer discussion as part of follow-up; as long as the instructor was using these additional activities to further discuss the original active learning activity, the additional activities were counted as follow-up. Although some instructors may ask rhetorical questions as part of a follow-up behavior, we decided to not try to distinguish between genuine and rhetorical questions. It was hard to do that consistently as observers, and we occasionally observed students treat rhetorical questions as genuine (Driessen et al., 2020). The end of the follow-up was marked by the instructor moving on from discussing the active learning activity, which was usually accompanied by advancing presentation slides.

Although CDOP and many other observation protocols have observers code the class continuously and record which codes are used in every two-minute intervals resulting in time distribution data (Kranzfelder et al., 2019), that strategy was not appropriate for FUDOP because follow-up behaviors occur sporadically throughout lecture, and their frequency varies depending on the number of active learning activities utilized by different instructors during a class session. In addition, we believed that there might be qualitative differences in how instructors followed up, so we were more concerned with which behaviors the instructors utilized for follow-up rather than how long these activities took. Therefore, we designed FUDOP to tally the number of times each FUDOP follow-up code occurred for every instructor follow-up segment after an active learning activity (Figure 1).

### Development of FUDOP codes

We engaged in a combination of deductive and inductive coding processes for the development of FUDOP codes (Fereday & Muir-Cochrane, 2006). This process was integral to answering our first research question, as it allowed us to use CDOP as a starting point while also incorporating unique instructor follow up behavior patterns we observed in large classes that were not observed in the more personal interaction setting explored by CDOP. Two researchers, X.C. and R.L., first used deductive coding by attempting to adapt the CDOP codes to recordings of six lectures taught by four different instructors from University A. At the same time, the researchers took note of any new behaviors they saw to inductively establish potential new codes. We refined the coding guide iteratively until X.C. and R.L. could establish a 100% consensus for each code from 30% of the class sessions from the University A data set. We then trained a third coder, M.K., and further tested the FUDOP coding scheme to code four courses taught by nine instructors from University B.

Finally, X.C. and M.K. applied the FUDOP coding scheme again in a new data set: two courses taught by six instructors at University C. No new codes were identified in this additional dataset, indicating that data saturation was reached (Fusch & Ness, 2015). Subsequent class sessions were coded by only 1 coder, and ambiguous cases were discussed by both coders. 12.5% of all the coded lectures were randomly selected for inter-rater reliability (IRR) measurement, and all codes had an IRR percent agreement ranging from 88.57% to 100% (Supplemental Table 2).

As we created the codes, we also characterized them as “instructor-centered” or “student-centered,” following Kranzfelder et al. (2019). If the code was derived from CDOP, we kept its classification in FUDOP. For our new codes, we designated the codes that involved multiple students or the whole class participating as “student-centered” and the codes that involved only one or no students participating as “instructor-centered.”

Instructor quotes included as examples in this manuscript (transcribed from video lecture recordings by X.C. manually) have been lightly edited for grammar and clarity in a way that enhances readability and retains the original intent of the instructor.

### Validation of FUDOP

Validity refers to the extent to which an instrument measures what it is intended to measure (Haynes et al., 1995). Specifically, content validity indicates that an instrument comprehensively represents components relevant to its subject of interest (in this case, instructor follow up behaviors). Relatedly, face validity indicates that users of an instrument view its elements as relevant to and consistent with their real-world experiences. To achieve both content and face validity for FUDOP, we engaged with both STEM instructors and researchers in science education from two universities.

Three of our participants in this process were both instructors and science education researchers, and three were primarily science education researchers. We provided the FUDOP coding scheme to these participants and facilitated two rounds of group discussions in which participants could collaboratively comment on code descriptions and example dialogues (Knekta et al., 2019). Authors X.C. and M.K. presented relevant background information and the FUDOP coding scheme with sample dialogues to the panel and asked the panelists: 1) to think about what they or other instructors might do after an active learning segment in their classes; 2) whether the code descriptions give a clear description of the new codes; 3) whether the brief example dialogues were sufficient for their understanding; and 4) if any of these codes applicable to their own follow-up behavior in the classroom. Then the panelists were presented with extended example dialogues, separate from the ones on the code description sheet, from a classroom and asked to think about which code or codes the instructor exhibited in their follow-up behaviors. Their feedback was collected to make clarifying changes and improve the usability of the FUDOP protocol.

### Analysis of follow up behavior patterns by instructor

To analyze follow up behavior patterns by instructors and identify similarities and differences within and between instructors to answer our second research question, we used R statistical software (R Core Team, 2019). To explore the differences between instructors in code usage, we used correspondence analysis, as this technique is appropriate for datasets containing multiple categorical variables (Sourial et al., 2010). We computed correspondence analysis using the CA() function from "FactoMineR" and "factoextra" packages (Kassambara & Mundt, 2020; Lê et al., 2008). We used biplots for the visual representation and identification of patterns and trends in the dataset. We plotted the correspondence biplot with fviz_ca_biplot(), displaying the top two dimensions that define the correspondence between the instructors and the FUDOP codes they utilize. The dimension contribution percentage represents the proportion of variance explained by each dimension. The cos2 (cosine square value) indicating how well each variable is represented along a specific dimension in the CA plot is also plotted with get_ca_row(), get_ca_col() functions, and corrplot from "corrplot" library (Wei et al., 2021). Color sets were from “RColorBrewer” library (Neuwirth, 2022). To compare patterns of instructor behaviors within themselves and between instructors, we employed Pearson’s chi-square test, as this test is appropriate for categorical data (Turhan, 2020). We computed Pearson’s chi-square values with chisq.test() in R to compare instructor behaviors within themselves for all FUDOP codes and between instructors for the top most frequent codes at each university.

## Results

### Development of the FUDOP codes

To address research question 1, we first analyzed the extent to which the CDOP codes could be observed during active learning follow-up. From the 17 total codes present in CDOP, we observed eight of them in the lectures: the four student-centered codes *Generative, Requesting, Checking-In,* and *Explaining,* and the four instructor-centered codes *Evaluating, Sharing, Linking,* and *Foreshadowing* (Kranzfelder et al., 2019).

Modifications were made to some CDOP codes to account for common follow-up behaviors. For example, the CDOP code *Explaining* occurs when a single student explains their thought process to other students, making it a one-sided monologue (Kranzfelder et al., 2019). In the mostly large lecture classes that were analyzed in this study, students were often asked to engage in conversations with one another to solve a problem together, resulting in a two-sided exchange. Because of this, we determined that *Neighbor Discussion* was a more accurate term than *Explaining* in the context of active learning follow-up behaviors. The CDOP code *Evaluating* is applied when an instructor acknowledges and reacts to a student’s response (Kranzfelder et al., 2019). We found that this descriptor was inadequate when applied to an instructor discussing the students’ collective responses to an active learning activity, so we modified it to *Student Answer-Oriented Summary.* The CDOP code *Sharing*, which is a general term used to describe any instance where the instructor provides information to students, was split into *General Summary* and *Hinting* in FUDOP to capture the specific nature of two distinct follow-up activities. We used *General Summary* to describe instances where instructors give a general explanation of the answer to an active learning activity and *Hinting* to describe instances where instructors give students guidance on how to approach the active learning activity without giving away the actual answer.

We also observed behaviors that were not described by CDOP. Four of them concerned different types of discourse: *Soliciting Individual Student’s Response*, *Announcing Answer, Delayed Follow-Up,* and *Prompting for Individual Thinking*. In addition, we observed instances where the active learning activity was not followed up by the instructor and there was therefore no discourse. We named this response *Moving on.* These nine CDOP-derived codes and five new codes were collated into a FUDOP coding guide (Supplemental Table 3).

When we solicited feedback on this original coding guide from a panel of STEM instructors and science education researchers, our panelists did not suggest adding or removing any codes. They did have suggestions on improving the code descriptions and names. The panelists informed us that it was hard to distinguish between *Generative* and *Requesting*, so we clarified that *Generative* can be answered in a closed-ended manner, often with a simple "yes" or "no" or a specific piece of information, whereas *Requesting* is prompting a more elaborate and detailed response, encouraging the student to express their thoughts and opinions. The panelists also had suggestions for the code *Neighbor Discussion.* We had observed that instructors sometimes speak during a peer discussion to encourage the students to continue to talk to each other, but we did not code that as a separate follow-up activity, since it represented a continuation of the same discourse move. The panelists suggested some small updates to the code description for clarity. They also suggested we change the name of the code to *Broadening conversation* to better match the format of the other codes as a behavior of the instructor. Additionally, panelists found it hard to distinguish between *Foreshadowing* vs *Delayed Follow-Up*, so we added more detailed explanations: *Foreshadowing* would mean the instructor performed the follow-up to the activity immediately afterward the activity but suggests the existence of ties to a future topic, whereas in *Delayed Follow-Up,* the follow-up activity itself is postponed until later. Finally, panelists also suggested minor edits of language for some code description and examples, which were incorporated into the FUDOP coding guide. The feedback from the expert panel on each code can also be found as part of Supplemental Table 3.

The panelists also brought up the issue of rhetorical questions, especially with the codes *Requesting* and *Checking-In.* They acknowledged that it would be hard to distinguish whether a question is rhetorical. We clarified that FUDOP does not attempt to distinguish between genuine and rhetorical questions, as an observer using FUDOP might find it difficult to tell the intention of the instructor.

### Explanation of Final FUDOP codes

The final FUDOP coding scheme that we developed to answer research question 1 consists of 14 codes: nine codes adapted from the CDOP framework, four codes that we developed through an iterative process of inductive coding, and one code for documenting when there is no discourse content (Table 2). Most of these codes can be categorized as student-centered or instructor-centered, with *Moving On* not falling into either of these categories.

**Table 2:**
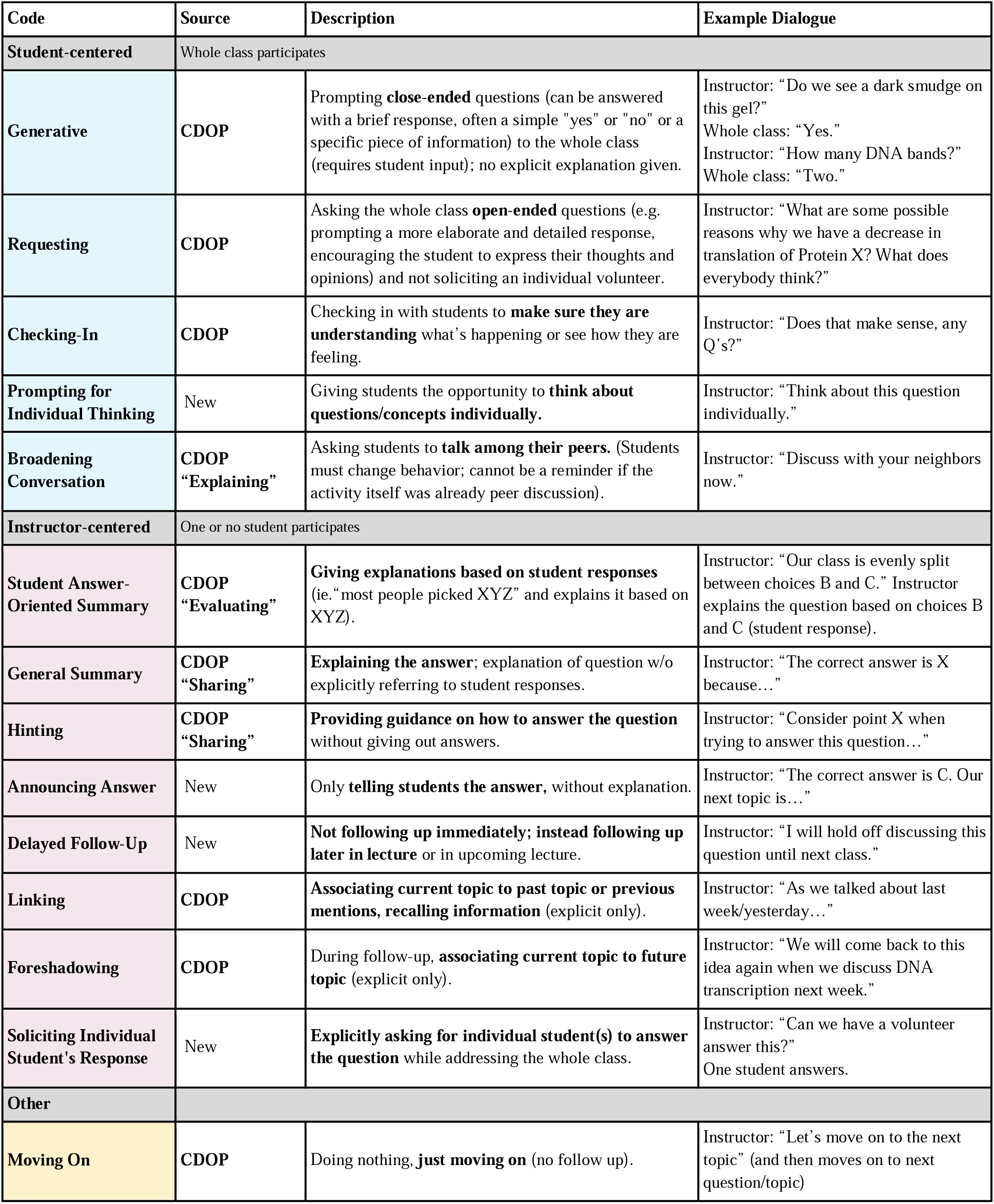
Finalized FUDOP Coding Scheme. FUDOP codes, separated into student-centered, instructor-centered, and other, with source, description, and example dialogue or situation.

Student-centered codes were defined as interactions where students in the whole class were given the opportunity to engage in the discussion. Five of the FUDOP codes are categorized as student-centered: *Generative, Requesting, Checking-In, Prompting for Individual Thinking, Broadening Conversation*. In three of these codes, the instructor asks questions without telling the students how to respond, and so any student who wanted to could respond. One such code is *Generative,* which involves instructors asking close-ended questions to the entire class without providing explicit explanations. An example of this code in action is the following:

Instructor: "Do we see a dark smudge on this gel?"
Class choral response: “Yes.”
Instructor: “How many DNA bands are there?”
Class choral response: “Two”.

In contrast, another code, *Requesting*, is characterized by instructors posing open-ended questions to the entire class, encouraging elaborate responses without soliciting individual volunteers. An instance of this is an instructor asking, "What are some possible reasons why we have a decrease in translation of Protein X? What does everybody think?" The *Checking-In* code captures instances where instructors are opening up the opportunity for students to lead their next move, which is routinely done to make sure that students understand the concept before moving on to the next concept. This code is exemplified by an instructor asking, "Does that make sense, any questions?" In the remaining two student-centered codes, the instructor directs all the students in the class to respond by engaging in active learning. The code *Prompting for Individual Thinking* involves instructors providing students with the opportunity to contemplate questions or concepts independently, as indicated by an instructor telling students to "Think about this question individually." The *Broadening Conversation* code entails instructors instructing students to engage and talk with their neighbors, fostering collaborative learning. An example is an instructor prompting, "Discuss with your neighbors now."

In comparison, instructor-centered codes describe interactions where the instructor holds the main voice in the classroom, giving students information that allows no room for the whole class of students to participate in the conversation. Eight FUDOP codes are categorized as instructor-centered: *Student Answer-Oriented Summary, General Summary, Hinting, Announcing Answer, Delayed Follow Up, Linking, Foreshadowing, Soliciting Individual Student’s Response*. In three of the codes, the instructor provides the answer. The *Announcing Answer* code entails instructors solely revealing the correct answer without offering an explanation, as demonstrated by an instructor stating, "The correct answer is C. Our next topic is…" With the code *General Summary*, instructors provide the answer but also explain it, such as an instructor stating, "The correct answer is D because…" In *General Summary, instructors do not* explicitly reference student responses. In contrast, with the code *Student Answer-Oriented Summary*, instructors provide explanations based on student responses, such as stating, "Our class is evenly split between choices B and C," and then explaining the question based on those choices. In two of the instructor-centered codes, the instructor explicitly does not provide the answer. With the *Hinting* code, instructors instead offer guidance and provide additional relevant details without providing direct answers, as exemplified by an instructor saying, "Consider the DNA sequences that can have specific regulatory functions when trying to answer this question." With the code *Delayed Follow-Up,* instructors do not provide the answer because they say they will follow-up later in the lecture or in subsequent lectures, as illustrated by an instructor saying, "I will come back to this question later." The codes *Linking* and *Foreshadowing* involve instructors associating the current topic with past or future topics, respectively.

In the case of *Linking*, instructors recall information from previous discussions, for example, an instructor stating, "As we talked about last week, …." Conversely, *Foreshadowing* captures instances where instructors connect the current topic to upcoming discussions, as seen in an instructor stating, "… And so, the correct answer here is an increase in transcription… We will continue discussing this topic in the next lecture." The last code in the instructor-centered category is *Soliciting Individual Student’s Response*. It involves instructors asking for individual students to answer questions by raising their hands. Although *Soliciting Individual Student’s Response* involves an individual student in the conversation during the follow-up, we still group this discourse under instructor-centered since the interaction is exclusive to the volunteer who got called on to answer the question. An instance would be an instructor asking, "Can we have a volunteer answer this?" This action excludes the majority of the class from the discussion and thus would not be considered student-centered.

Finally, the *Moving On* code, which is neither instructor-centered nor student-centered, describes situations where there is no discourse. Instructors take no further action and proceed to the next topic without additional follow-up. An example is an instructor stating, "Let’s move on to the next topic," without providing the answer or any further talk about the question, and immediately transitioning to the subsequent question or topic.

As an example, to demonstrate the process of FUDOP coding, we will analyze an example of how an instructor followed up after a single clicker question on regulation of gene expression. The instructor had given a clicker question and allocated some time for students to discuss before the instructor regrouped the class.

Instructor: “The class was split 50-50. Can those who answered B give me a reason? It’s good that so many people answered B. Why did you pick B?
An individual student raises their hand, the instructor calls on them, and the student gives an answer.
Instructor: “So let me see if I get your reason. You think the enhancer may or may not be needed for transcription. Is that correct? How about we put it in a different way? What does the ‘enhancer’ mean-to enhance something? So if we are talking about transcription, the only meaning for “enhancer” is to enhance transcription. So enhancer A is a DNA sequence, DNA element, usually 68 to 80 nucleotides long. Very Short. They are there to enhance transcription. How can a piece of DNA that enhances transcription occur at the promoter? That’s your question, right? You think well, because the promoter is required for transcription, enhancement may or may not be required for transcription. That is correct. Remember the CAP protein we talked about the last time-you got CAP binding to the binding site, the promoter is downstream, then you push it on the polymerase to activate transcription. So you need to bind to something in order to activate transcription, right? An activator. So, an activator is a transcription factor, a positive regulator of transcription-that means it activates transcription. It activates transcription. If you don’t have an enhancer or the enhancer cannot bind to the activator,
what will happen to transcription?
Class choral response: “Decrease.”
Instructor: “Does that make sense? Good?”

The first discourse move the instructor made was to call for an individual who picked choice B to answer the question, by asking “Can those who answered B give me a reason?” and then calling on an individual student who raised their hand. This interaction would be coded under the FUDOP code *Soliciting Individual Student’s Response*. Next, the instructor explained a biology concept by linking it to a previously discussed topic when they said, “Remember the CAP protein we talked about the last time?” This discourse move would be coded under the FUDOP code *Linking*. After that, the instructor asked a question of the whole class when they queried, “If you don’t have an enhancer or the enhancer cannot bind to the activator, what will happen to transcription?” This discourse move falls under the code *Generative*. The final discourse move the instructor made is checking in with the class by asking, “Does that make sense?” This move would be marked as “Checking-In”. More examples including this one coded with FUDOP are shown in Supplemental Figure 1.

### Instructors show distinct patterns of follow-up behaviors

To answer research question 2, about whether instructors differed in their follow-up behaviors, we used FUDOP to analyze and measure instructor behaviors. We hypothesized that FUDOP would allow us to see differences between instructors, as this new protocol has the potential to reveal interesting patterns that were previously unquantifiable and might be important for modulating the effects of active learning. To examine the overall pattern of instructor behavior, we generated 100% stacked bar graphs of follow-up codes in all sessions per instructor (Figure 2). Visual inspection revealed that although there are commonly occurring codes like *General Summary* that are often used by the majority of the instructors (N=18), instructors appeared to differ from each other in the overall pattern of FUDOP code utilization. This observation was confirmed with a chi-square test (p<0.001). Some university-specific patterns also emerged, such as the fact that *Soliciting Individual Student’s Response* exclusively occurred at universities B and C.

To test the nascent hypothesis that individual instructors use distinct follow-up behaviors, we performed a correspondence analysis of the relationship between instructors and follow-up discourse behaviors. We chose to perform correspondence analysis to visualize the association between the two categorical variables “instructor” and “FUDOP codes” because it allowed us to determine the strength and direction of associations between categories. According to the correspondence analysis, there were three main dimensions: Dim1 (29.9% of variance), Dim2 (22.2% of variance), which together contributed to two thirds (67%) of the variance observed in our dataset (Figure 3A, Supplemental Figure 2). The quality of representation of the variables in each dimension can be quantified using the square cosine (cos2); a high cos2 value indicates how well each category or variable is represented along a specific dimension. Analysis of cos2 values indicated that Dim1 was largely defined by the dichotomy between the codes *Individual Student’s Response* and *Moving On* as well as instructors A2 and C2 (Figure 3A, which may reflect an instructor’s willingness to engage in follow-up behavior at all. Dim2 was defined by the dichotomy between *Announcing Answer* and *General Summary* (Figure 3B), which may reflect an instructor’s preference about simply announcing the answer as opposed to engaging in more in-depth follow-up activities. Dim3 was solely defined by the presence or absence of the *Checking-In* code (Figure 3B), which may reflect instructor preference on whether to check in with students.

**Figure 3:**
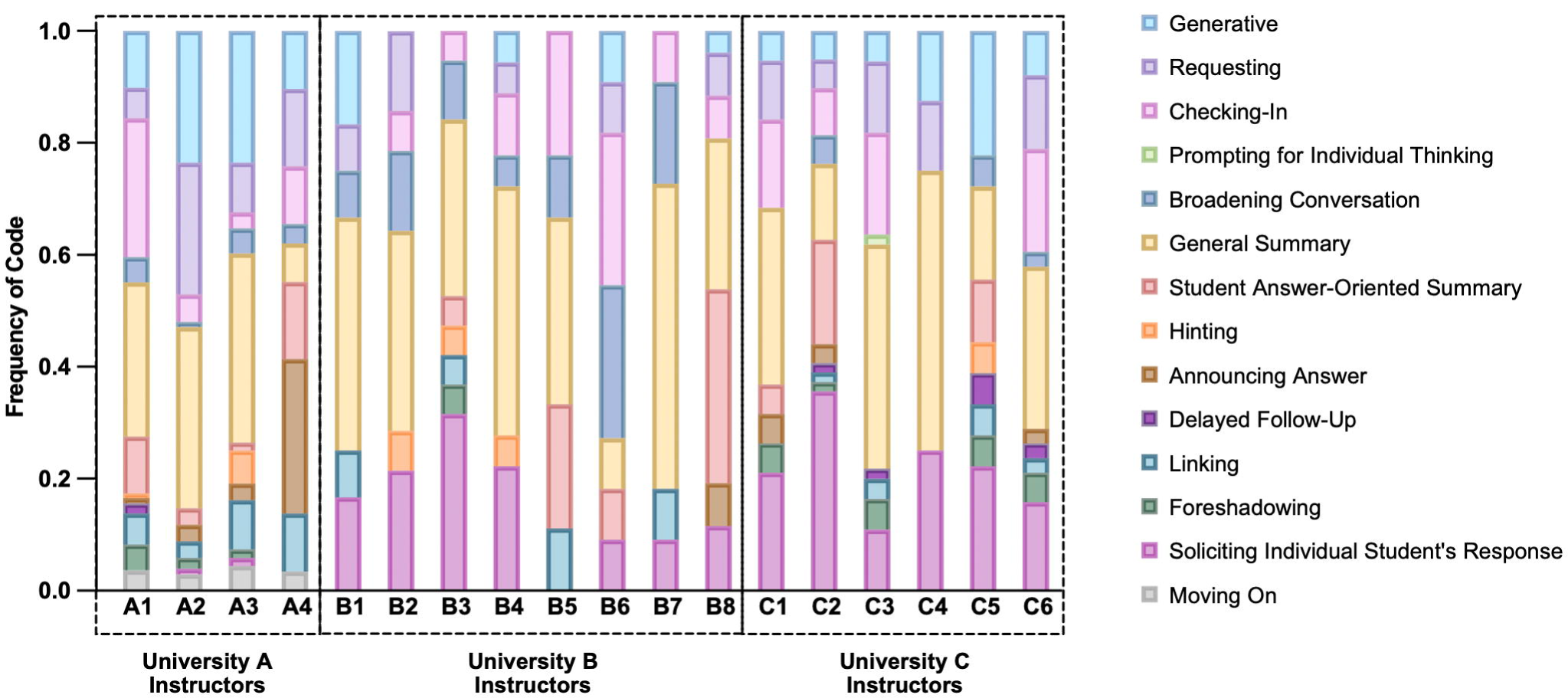
Instructors display distinct patterns of follow-up methods from each other. 100% stacked frequency bar graphs of follow-up codes in all sessions.

Analysis of the correspondence plot pointed to there being significant differences between instructors. We observed that different instructors appeared near various codes (Figure 3A). For instance, instructor A2 and A3 appeared near *Generative* and *Requesting*, which both involve the instructor posing questions to the students. That contrasted with instructor B8, who distinctly appeared on the plot near *Student Answer-Oriented Summary*. Another significant contrast was observed on Dim1 with instructor B3 appearing near *Soliciting Individual Student’s Response*, compared to instructor A4 appearing on the other end of the plot near *Announcing Answer*. Interestingly, instructors from University A were all located on one side of the biplot, whereas instructors from the other two universities were closer together on the other end of the biplot, indicating that instructors from University A exhibited more similar follow-up behaviors to each other than to instructors from other institutions. The chi-square of independence between the two variables indicates that there are significant differences in the proportion of various codes used by different instructors (X^2^ (221, N = 625) = 444.162, p = 4.04x10^-17^).

To further address research question 2, we not only wanted to examine whether instructors had different patterns of follow-up behaviors but also whether different instructors spent different amounts of time in absolute terms performing various follow-up behaviors. Therefore, we examined how often each code was used per minute of class time and whether we could detect significant differences between instructors in code usage (Figure 5, Supplemental Table 4). Because we saw university-specific differences in code usage, we graphed the instructors at each university separately (Figure 5A, C, E) and compared them only to instructors and their own university (Figure 5B, D, F). While Figure 4 shows the comparisons for the most common five codes at each university to highlight which codes could differ the most between instructors, Supplemental Table 4 shows the same comparison for all codes. At University A, usage of *Requesting* and *Checking-In* was significantly different between instructors (Figure 5B). In contrast, at University B, only usage of the *Student Answer-Oriented Summary* code demonstrated a significant difference between the instructors (Figure 5D), while at University C, none of the five most frequently used codes at this university showed a significant difference between the instructors, indicating higher uniformness in follow up behaviors (Figure 5F).

**Figure 4:**
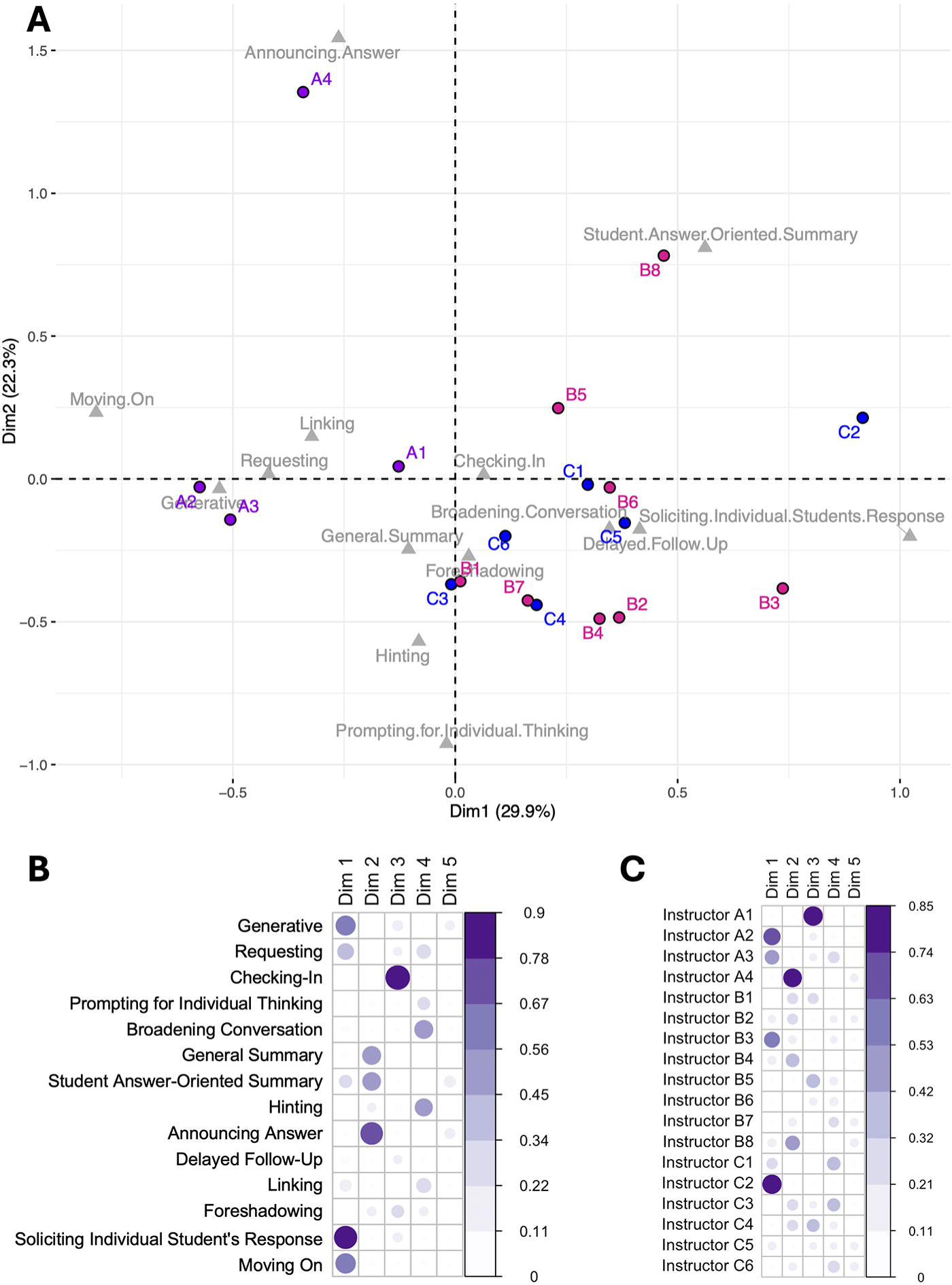
Different instructors had patterns of behavior that cluster with different sets of codes. (A) Correspondence analysis biplot of the relationship between instructors and follow-up code discourse behavior. (X-squared of independence between the two variables is equal to 444.162, p-value = 4.04x10^-17^) (B) Cosine squared (cos2) values for the association between follow-up code discourse behaviors and the first five dimensions from the correspondence analysis. Higher cos2 values indicate stronger associations with the respective dimension. (C) cos2 values for the association between individual instructors and the first five dimensions from the correspondence analysis.

**Figure 5:**
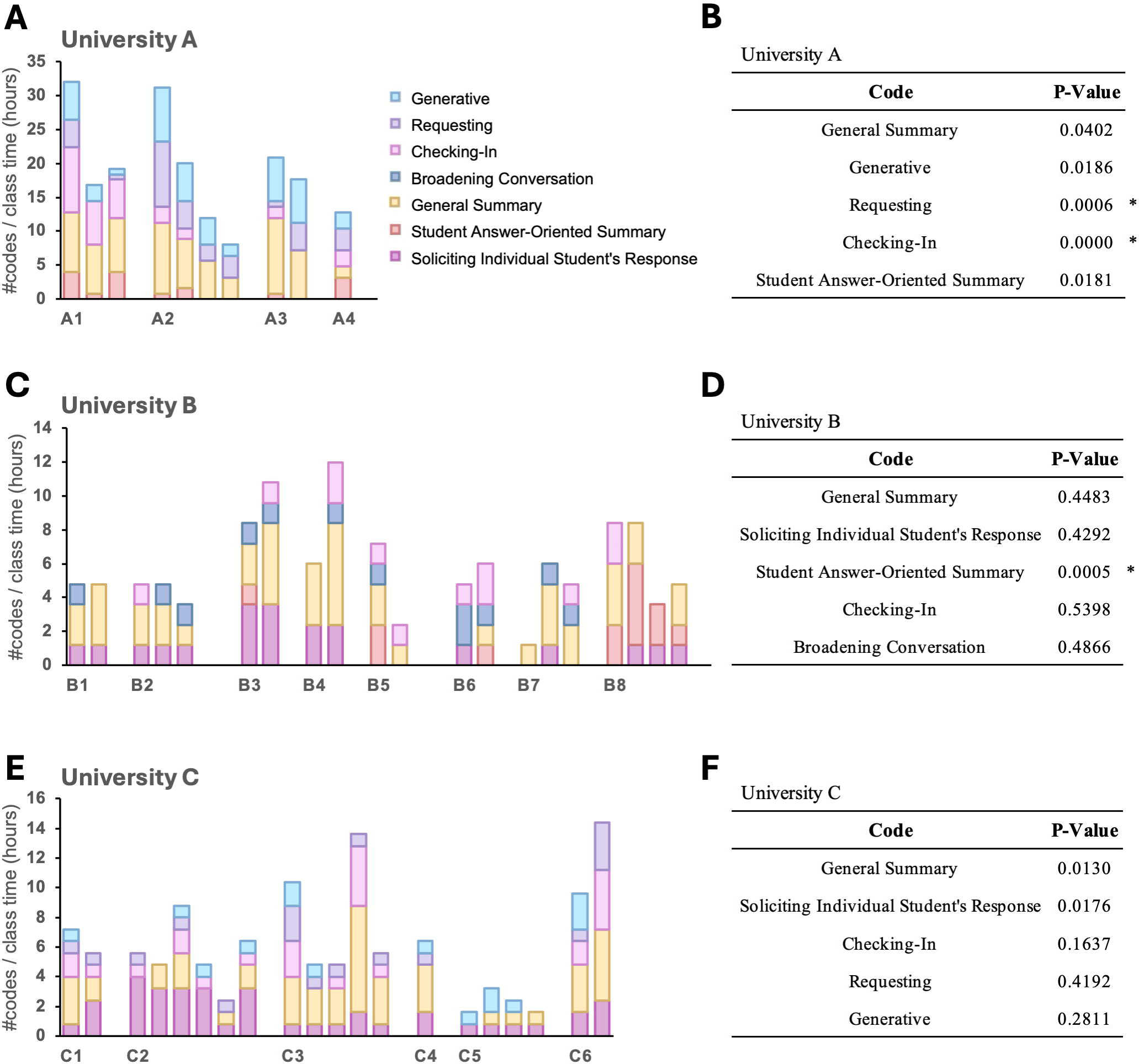
Five most frequently used codes per institution and whether their use differed by instructor. Stacked bar graphs (A, C, E) and Pearson’s chi-square test p-values (B, D, F) of the top 5 codes used at each institution. All the class sessions analyzed for each instructor are plotted, so one column is one class session. For B, D, F, * designates significance after correction for multiple comparisons (p < 0.003).

Because follow-up behaviors can only be performed after active learning activities and varied in how often they performed those (Supplemental Figure 3A), we also examined how often each code was used per active learning activity (Supplemental Figure 3B). Using more codes per active learning activity may indicate richer or more diverse discourse between the instructor and the class. Most instructors, but not all, used more than one follow-up activity on average after a single active learning activity (Supplemental Figure 3B). Also, some instructors used different follow-up activities for every active learning activity, but others tend to use the same ones each time. For instance, instructor A4 only performed approximately one follow-up behavior per activity, but that behavior could be any of five different FUDOP codes. In contrast, instructor B2 did 2-3 follow-up behaviors per activity, but they varied between only four different FUDOP codes (Supplemental Figure 3B).

### Instructors tended to behave consistently in different class sessions

When addressing research question 2, we might have been able to discern differences in FUDOP code usage between instructors either because instructors had roughly consistent patterns of follow-up usage that they use when teaching introductory biology or because instructors used different follow-up behaviors when teaching different topics. Therefore, we set out to answer an additional research question: do individual instructors use consistent discourse behaviors across different class sessions? To answer this question, we used FUDOP to analyze whether each instructor behaved roughly consistently in different class sessions, which address different material.

A heatmap of codes by all class sessions taught by each instructor showed similar patterns of code frequency across class sessions within each instructor themselves (Figure 5A). For instance, instructors A1 and B7 were fairly different from each other but were both internally consistent: instructor A1 consistently did a lot of *Checking-In*, while instructor B7 consistently did a lot of *General Summary* (Figure 5A). We performed a chi-square test with adjustment for multiple comparisons for each instructor with multiple class sessions to see whether each instructor behaved consistently in different class sessions. There were no significant differences in code usage by class session for class sessions taught by the same instructor, which supported the idea that each instructor behaved consistently in different class sessions (Figure 5B).

## Discussion

### Our findings

An instructor’s response to students after an active learning activity can be conceptualized as a form of discourse. Therefore, we created and implemented an adapted version of the CDOP coding scheme, which we have termed the Follow-Up Discourse Observation Protocol (FUDOP), to categorize instructors’ follow-up discourse move behaviors after an active learning activity (Kranzfelder et al., 2019). We showed that we could consistently apply our new coding scheme to instructors across three different universities with diverse institutional characteristics. Using FUDOP, we found that we could identify significant differences between instructors in their follow-up behaviors, even between instructors in the same institution, but that the instructors were roughly consistent in which follow-up behaviors they exhibited in different class sessions.

### Importance of discourse in the college STEM classroom

Previous research strongly suggests that the type of discourse moves chosen by an instructor matters. Certain discourse patterns can better engage students or prompt students to use higher-order thinking (Alkhouri et al., 2021; Bae et al., 2021; Chin, 2006; Van Booven, 2015; Zhang, 2009). Either of these outcomes might lead to more student learning.

As more instructors of college STEM courses adopt active learning-based pedagogies, there are more opportunities to use diverse discourse moves (Kranzfelder et al., 2019; Stains et al., 2018). In the traditional large lecture-based STEM college classroom, the discourse patterns used were largely limited to those that were authoritative and non-interactive, involving the instructor lecturing (Mortimer & Scott, 2003). Even if the instructor took student questions or solicited individual student responses, the typical student remained a passive recipient of information and was not involved in interaction and the co-construction of knowledge (Conner et al., 2024). With active learning pedagogies, however, instructors have many more options for how they can interact with and engage students. For example, they can interact with students as they are completing activities in class (Kranzfelder et al., 2019). They can also interact with the students as a whole during active learning activities, as students complete activities and share their answers and outcomes with the instructor (Hattie & Timperley, 2007; Wood et al., 2018).

Measuring and quantifying the discourse patterns found in active learning classrooms may have important implications for better understanding and improving outcomes associated with active learning classrooms. Even though active learning has been shown to generally increase student learning, individual studies show effects of various sizes, with some studies showing no benefit over lecturing (Freeman et al., 2014). It is not clear why that is, but it is known that, even with the efforts put into adopting active-learning-based teaching strategies across universities, there is still significant variability in how instructors deliver these strategies in classrooms (Stains et al., 2018). Although there have been numerous studies aiming to identify the most effective teaching methods, we still do not have a mechanistic understanding of how variability in instructors’ implementation strategies impact student learning (Martella et al., 2024; Streveler & Menekse, 2017). In order to gain this mechanistic understanding, researchers must first dissect the different cognitive processes and dialogic interactions that make up an activity learning activity and figure out their relative contributions (Vickrey et al., 2015). One of these components is instructor follow-up, as this is an important way that instructors interact and give feedback to their students in a large classroom (Hattie & Timperley, 2007; Wood et al., 2018). Therefore, we believe that FUDOP could be used to identify differences in follow-up behaviors between instructors, allowing future work to investigate how differences in these behaviors could lead to different student outcomes.

### Relationship between FUDOP codes and previous research on discourse

The FUDOP codes grew out of previous research on discourse. According to Mortimer and Scott, discourse moves can be categorized into two axes of approach: dialogic-authoritative and interactive-non-interactive (Mortimer & Scott, 2003). The FUDOP codes that describe follow-up behaviors can also be placed into this theoretical framework (Figure 6). The codes *Requesting, Broadening Conversation,* and *Soliciting Individual Student’s Response* can be characterized as dialogic, since student perspectives are being discussed, and interactive, since the students are actively interacting with each other or the instructor. In contrast, the *Generative* and *Checking-In* codes fell into the category of authoritative and interactive because there is still interaction between the instructor and the students (interactive), but the ideas discussed in the classroom are solely the instructor’s (authoritative). In the dialogic-noninteractive category, we have *Prompting for Individual Thinking* and *Student Answer-Oriented Summary.* These codes allow for student input but do not involve interaction between the instructor and students. We then categorized *General Summary, Hinting, Announcing Answer, Delayed Follow-Up, Linking,* and *Foreshadowing* as authoritative-non-interactive because they only feature the instructor’s perspective (authoritative) and do not involve student interaction (non-interactive). Defining the discourse characteristics of instructor follow up behaviors in this way can provide insight into how students learn in different active learning classrooms, as evidenced by previous work describing the various impacts of these discourse moves (Alkhouri et al., 2021).

**Figure 6:**
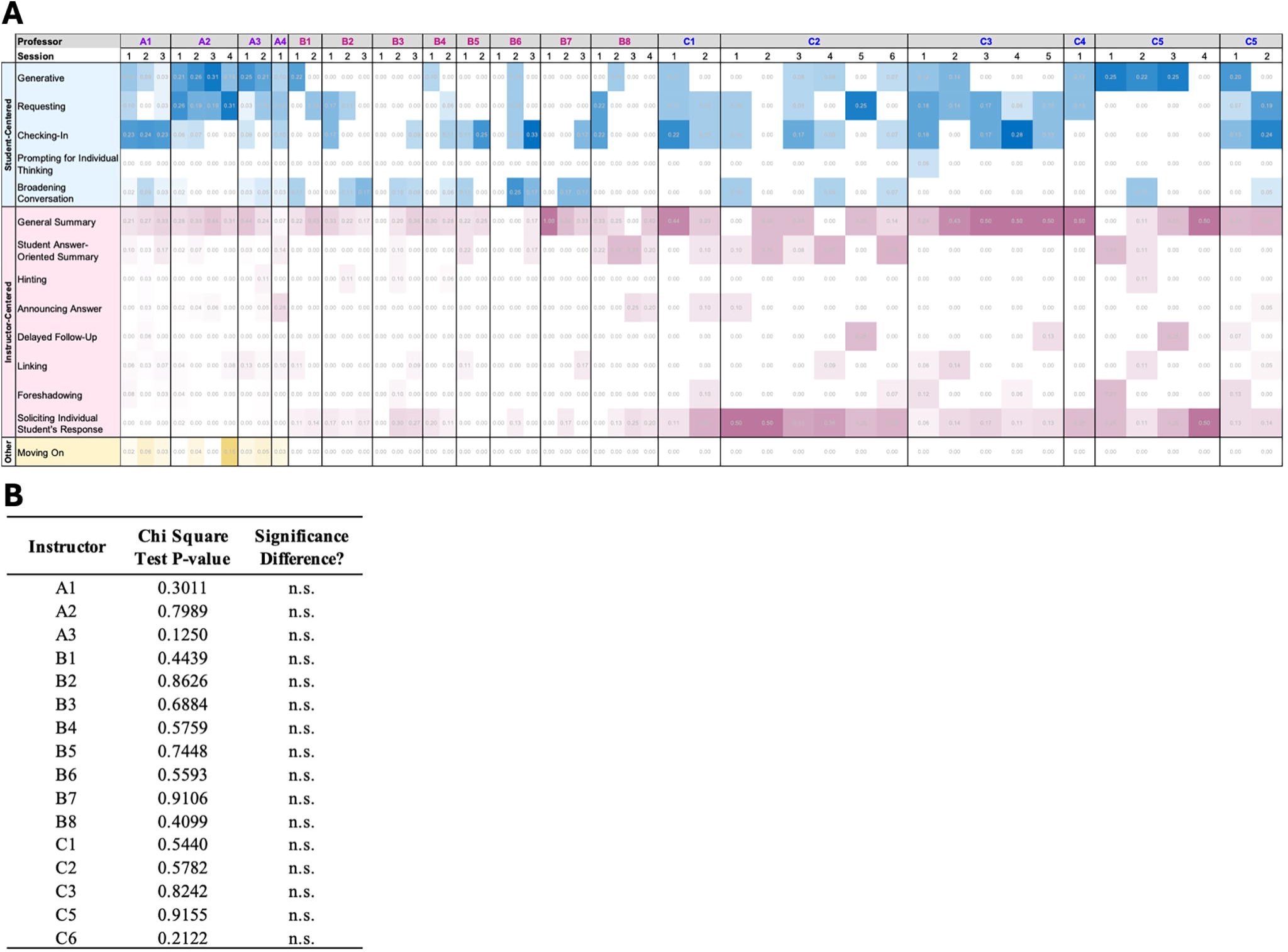
Frequencies of codes at each institution by instructor and class session. (A) Heat map of student-centered codes in blue, instructor-centered codes in pink, and others in light yellow. Each column is one class session, and class sessions taught by the same instructor are boxed together. (B) Pearson’s chi-square test p-values for instructors with multiple class sessions.

**Figure 7:**
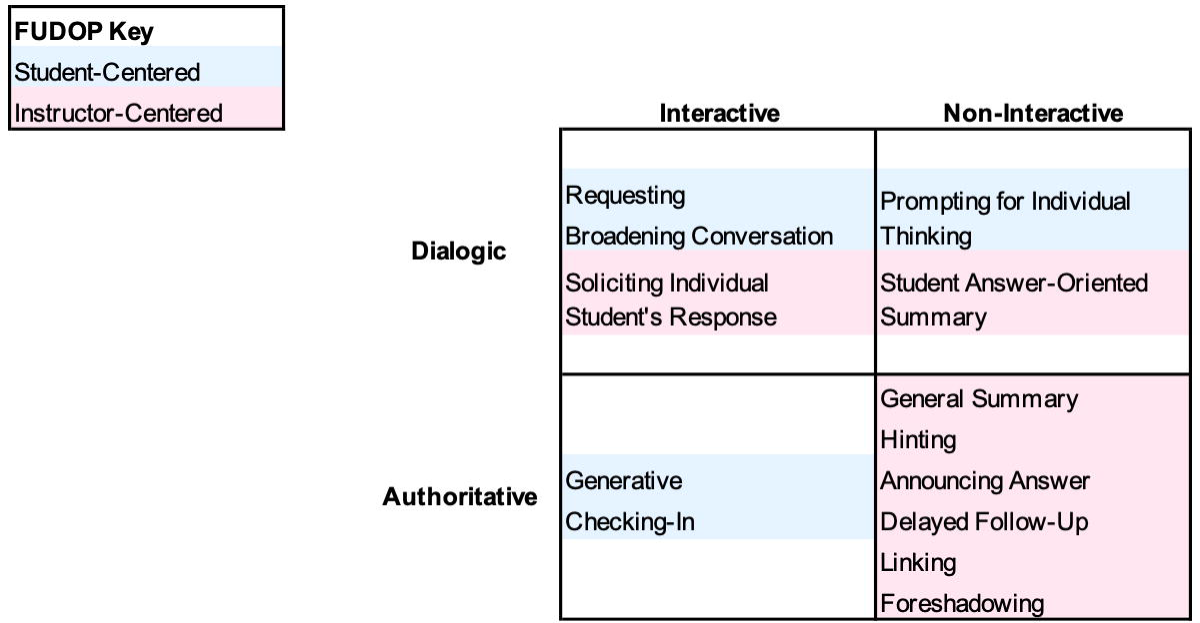
FUDOP codes along the two axes of approach (Mortimer & Scott)

Following CDOP, we also categorized our codes as either “student-centered” or “instructor-centered” (Kranzfelder et al., 2019). “Student-centered” codes were ones in which a majority of students could, at least in theory, participate, while “instructor-centered” codes were ones in which only the instructor or only a handful of students, at most, could participate. These categories mostly but not entirely aligned with the interactive-non-interactive approach. *Solicit Individual Students Response* was interactive because at least one student was interacting, but we did not designate it as “student-centered” because in a large college class, the vast majority of the students are merely passively listening when one or a few students are responding. Similarly, *Individual Thinking* was non-interactive because thinking silently is not interactive, but we designated it as “student-centered” because all the students in the class are being exhorted to think. Further research is necessary to determine the cognitive impact of student-versus instructor-centered follow-up behaviors.

### Variability in instructor follow-up strategies

We found that although instructors behaved consistently from session to session (Figure 5), many instructors used codes in a way that were significantly different from other instructors (Figure 3). It is possible that instructors who teach at different universities tend to show different types of follow-up behaviors because these behaviors are influenced by factors that can vary by university or department. These factors could include administrative policies, training, support for innovative pedagogy, classroom context, and student factors, which may affect the use of evidence-based teaching in general (Cavanagh et al., 2018; Lau et al., 2024; Owens et al., 2018; Stains et al., 2018). However, differences were even seen among instructors who taught at the same institution (Figure 4). The differences seen between instructors in University A is particularly striking because all instructors at University A were given the same materials for the common course they all taught, including the same slides and the same clicker questions (Figure 4A, B). This means they had the same number and type of opportunities to do active-learning follow-up up with the students, and they still showed a significant difference in their follow-up methods when compared to each other.

It is not known what the source of this variability between instructors is. The fact that instructors at University A used significantly different follow-up strategies suggests that the variability does not solely arise from differences in the topics taught or the questions used, as all of these instructors used the same slides and clicker questions. Rather, the variability might arise from differences in their experiences (or lack of experience) learning in active-learning classrooms. It could also arise from training experiences. Although some forms of active learning, like peer instruction, clearly prescribe certain forms of follow-up (Mazur, 1997), not all types of active learning do, and in any case even when there are clear guidelines, like with peer instruction, not all instructors follow them faithfully (Stains & Vickrey, 2017). Therefore, currently, there is ample room for instructors to use their judgment and experience to choose which follow-up strategies to use in particular situations. The variability we found also suggests that there could be opportunities for targeted teaching professional development that specifically addresses how to select follow-up strategies.

Differences between instructors in their approaches to follow-up interactions could also potentially be attributed to variations in their conceptions of teaching and learning (Ertmer & Newby, 2013; Rozhenkova et al., 2023; Woolley et al., 1999). Instructors with behaviorist orientations may prioritize correction and reinforcement, aiming to inform student misunderstandings directly (Schunk, 2012). For instance, they might favor FUDOP codes, like *Announcing Answer* and *General Summary*. In contrast, those with constructivist orientations may choose instructor behaviors that focus on facilitating student sense-making and constructing knowledge collaboratively (Schunk, 2012). In this case, FUDOP codes *Requesting*, *Prompting for Individual Thinking*, *Broadening Conversation*, and *Hinting* may be consistent with constructivist learning theories as they are behaviors that direct students to come up with their own conception of the topic. If differences between instructors arise mainly from conceptions of teaching and learning, the fact that instructors are mostly consistent from session to session also makes sense. Consistency within instructors may arise from the stability of their underlying conceptions over time. Instructors tend to develop and refine their pedagogical beliefs and practices based on their experiences, professional development opportunities, and disciplinary norms (Luft et al., 2003; Tondeur et al., 2017). As a result, instructors may exhibit consistency in their follow-up discourse patterns within their own teaching contexts, reflecting the coherence of their instructional philosophies and approaches. Further research could explore whether and how conceptions of teaching and learning predict follow-up behaviors.

### Limitations and Future Directions

There are many limitations to our work. First, FUDOP was only applied to three universities. Although the institutions were diverse and we reached code saturation in the second university we examined, we cannot rule out the possibility that there are other follow-up strategies besides the ones we observed. In particular, there might be codes that are very rarely used in biology classrooms but are more commonly used in other STEM disciplines. Future studies might uncover additional follow up strategies by deploying FUDOP in different educational contexts. Second, FUDOP only captures a small part of the potential discourse interactions in the classroom, specifically follow-up after active learning. Other observation protocols would need to be used to capture other aspects of the classroom, such as discourse between the instructor and individual students or small groups; the general type of activities used by the instructor, and what practices are used out-of-class to help students learn. A future study combining these protocols to create a more holistic understanding of classroom instructional practices might be useful. Finally, FUDOP does not capture the instructor’s own perspectives, which may miss essential insights into their intentions and teaching goals. Future qualitative studies should investigate instructor intent to better understand why different instructors utilize various follow up behaviors.

### Practical applications of FUDOP

FUDOP is of potential interest to both researchers and educators. FUDOP can be used by researchers as a tool to investigate questions related to active learning follow-up. For example, future research could explore how the cognitive level of the active learning activities influence which discourse moves are used to follow up. Importantly, FUDOP could also be used to study the relative effectiveness of various follow-up practices, giving educational researchers valuable insights into the dynamics of the classroom to find effective pedagogical practices and evidence-based teaching methods.

Furthermore, FUDOP establishes a unique fingerprint of recurring codes that are specific to the instructor, which could provide researchers a better understanding of teaching practices for professional development. FUDOP can also be used by educators. Studies show that instructors commonly mis-estimate how often they use various teaching techniques (Ebert-May et al., 2011; Sheridan & Smith, 2020), but observation protocol data can help instructors better understand and improve their own teaching (Pianta & Hamre, 2009; Reisner et al., 2020). FUDOP allows instructors to review their discourse and interactions with students, which could highlight areas for improvement in their teaching skills and thereby foster more engaging and effective classroom environments. For example, an instructor who prides themselves on extensively using student-centered learning techniques might see with FUDOP that, while they do frequently use active learning activities like clicker questions, they primarily use instructor-centered follow-up activities like *general summary* and *soliciting individual student’s responses.* That insight might prod them to try more student-centered techniques like *broadening conversation* or *prompting for individual thinking*. These types of insights have been anecdotally noted when using other observation protocols like COPUS for instructor professional development (Reisner et al., 2020).

## Conclusion

In very large classes, like most of the ones studied here, where opportunities for one-on-one interaction with the instructor are very limited, instructor feedback after active learning may be the only dialogic experience many students have with the instructor. There is a compelling need to study instructor follow-up more extensively: this essential aspect of the teaching process is often overlooked in educational research, despite its meaningful role in enhancing learning (Smith et al., 2011). The follow-up adapted codes in the FUDOP coding scheme allow for the structured observation and categorization of the implementation behaviors of instructors during this important part of active learning. FUDOP can be used as a tool for researchers and educators to better understand what follow-up behaviors are being used and, in the future, tie those behaviors to educational outcomes.

## Supporting information

All supplemental materials

## Acknowledgements

We would like to thank the UCSD and UCLA biology education research communities for helpful discussions and to all of the instructors who participated in this study. We would also like to thank the UC Merced SATAL interns for help with data collection. Collection of the data was supported by Howard Hughes Medical Institute award no. GT11066.

## References

Alkhouri, J. S., Donham, C., Pusey, T. S., Signorini, A., Stivers, A. H., & Kranzfelder, P. (2021). Look Who’s Talking: Teaching and Discourse Practices across Discipline, Position, Experience, and Class Size in STEM College Classrooms. BioScience, 71(10), 1063– 1078. 10.1093/biosci/biab077

Auerbach, A. J., & Schussler, E. E. (2017). Curriculum Alignment with Vision and Change Improves Student Scientific Literacy. CBE—Life Sciences Education, 16(2), Article 2. 10.1187/cbe.16-04-0160

Bae, C. L., Mills, D. C., Zhang, F., Sealy, M., Cabrera, L., & Sea, M. (2021). A Systematic Review of Science Discourse in K–12 Urban Classrooms in the United States: Accounting for Individual, Collective, and Contextual Factors. Review of Educational Research, 91(6), Article 6. 10.3102/00346543211042415

Blanton, M. L. (2002). Using an Undergraduate Geometry Course to Challenge Pre-service Teachers’ Notions of Discourse. Journal of Mathematics Teacher Education, 5(2), 117–152. 10.1023/A:1015813514009

Boud, D., & Molloy, E. (2013). Rethinking models of feedback for learning: The challenge of design. Assessment & Evaluation in Higher Education, 38(6), Article 6. 10.1080/02602938.2012.691462

Boveda, M., & Annamma, S. A. (2023). Beyond Making a Statement: An Intersectional Framing of the Power and Possibilities of Positioning. Educational Researcher, 52(5), Article 5. 10.3102/0013189X231167149

Cavanagh, A. J., Chen, X., Bathgate, M., Frederick, J., Hanauer, D. I., & Graham, M. J. (2018). Trust, Growth Mindset, and Student Commitment to Active Learning in a College Science Course. CBE—Life Sciences Education, 17(1), Article 1. 10.1187/cbe.17-06-0107

Chin, C. (2006). Classroom Interaction in Science: Teacher questioning and feedback to students’ responses. International Journal of Science Education, 28(11), Article 11. 10.1080/09500690600621100

Conner, J., Mitra, D. L., Holquist, S. E., Rosado, E., Wilson, C., & Wright, N. L. (2024). The pedagogical foundations of student voice practices: The role of relationships, differentiation, and choice in supporting student voice practices in high school classrooms. Teaching and Teacher Education, 142, 104540. 10.1016/j.tate.2024.104540

Dewsbury, B. M., Swanson, H. J., Moseman-Valtierra, S., & Caulkins, J. (2022). Inclusive and active pedagogies reduce academic outcome gaps and improve long-term performance. PLOS ONE, 17(6), Article 6. 10.1371/journal.pone.0268620

Donham, C., & Andrews, T. C. (2023). “What’s Your Thinking behind That?” Exploring Why Biology Instructors Use Classroom Discourse. Journal of Microbiology & Biology Education, 24(2), e00132–22. 10.1128/jmbe.00132-22

Driessen, E. P., Knight, J. K., Smith, M. K., & Ballen, C. J. (2020). Demystifying the Meaning of Active Learning in Postsecondary Biology Education. CBE—Life Sciences Education, 19(4), ar52. 10.1187/cbe.20-04-0068

Ebert-May, D., Derting, T. L., Hodder, J., Momsen, J. L., Long, T., & Jardeleza, S. (2011). What We Say Is Not What We Do: Effective Evaluation of Faculty Professional Development Programs. BioScience, 61(7), 550–558.

Ertmer, P. A., & Newby, T. J. (2013). Behaviorism, Cognitivism, Constructivism: Comparing Critical Features From an Instructional Design Perspective. Performance Improvement Quarterly, 26(2), Article 2. 10.1002/piq.21143

Fereday, J., & Muir-Cochrane, E. (2006). Demonstrating Rigor Using Thematic Analysis: A Hybrid Approach of Inductive and Deductive Coding and Theme Development. International Journal of Qualitative Methods, 5(1), 80–92. 10.1177/160940690600500107

Ford, M. J., & Wargo, B. M. (2012). Dialogic framing of scientific content for conceptual and epistemic understanding. Science Education, 96(3), Article 3. 10.1002/sce.20482

Freeman, S., Eddy, S. L., McDonough, M., Smith, M. K., Okoroafor, N., Jordt, H., & Wenderoth, M. P. (2014). Active learning increases student performance in science, engineering, and mathematics. Proceedings of the National Academy of Sciences, 111(23), Article 23. 10.1073/pnas.1319030111

Fusch, P., & Ness, L. (2015). Are We There Yet? Data Saturation in Qualitative Research. The Qualitative Report, 20(9), Article 9. 10.46743/2160-3715/2015.2281

Hattie, J., & Timperley, H. (2007). The Power of Feedback. Review of Educational Research, 77(1), Article 1. 10.3102/003465430298487

Haynes, S. N., Richard, D. C. S., & Kubany, E. S. (1995). Content validity in psychological assessment: A functional approach to concepts and methods. Psychological Assessment, 7(3), Article 3. 10.1037/1040-3590.7.3.238

Holmes, A. G. D. (2020). Researcher Positionality—A Consideration of Its Influence and Place in Qualitative Research—A New Researcher Guide. Shanlax International Journal of Education, 8(4), Article 4.

Kassambara, A., & Mundt, F. (2020). factoextra: Extract and Visualize the Results of Multivariate Data Analyses (Version 1.0.7) [Computer software]. https://cran.r-project.org/web/packages/factoextra/index.html

Knekta, E., Runyon, C., & Eddy, S. (2019). One Size Doesn’t Fit All: Using Factor Analysis to Gather Validity Evidence When Using Surveys in Your Research. CBE—Life Sciences Education, 18(1), rm1. 10.1187/cbe.18-04-0064

Kranzfelder, P., Bankers-Fulbright, J. L., García-Ojeda, M. E., Melloy, M., Mohammed, S., & Warfa, A. R. M. (2019). The Classroom Discourse Observation Protocol (CDOP): A quantitative method for characterizing teacher discourse moves in undergraduate STEM learning environments. PLoS ONE, 14(7), 1–20. 10.1371/journal.pone.0219019

Lau, A. C., Henderson, C., Stains, M., Dancy, M., Merino, C., Apkarian, N., Raker, J. R., & Johnson, E. (2024). Characteristics of departments with high-use of active learning in introductory STEM courses: Implications for departmental transformation. International Journal of STEM Education, 11(1), Article 1. 10.1186/s40594-024-00470-x

Lê, S., Josse, J., & Husson, F. (2008). FactoMineR: An R Package for Multivariate Analysis. Journal of Statistical Software, 25, 1–18. 10.18637/jss.v025.i01

Lemke, J. L. (2001). Articulating communities: Sociocultural perspectives on science education. Journal of Research in Science Teaching, 38(3), 296–316. 10.1002/1098-2736(200103)38:3<296::AID-TEA1007>3.0.CO;2-R

Lombardi, D., Shipley, T. F., Bailey, J. M., Bretones, P. S., Prather, E. E., Ballen, C. J., Knight, J. K., Smith, M. K., Stowe, R. L., Cooper, M. M., Prince, M., Atit, K., Uttal, D. H., LaDue, N. D., McNeal, P. M., Ryker, K., St. John, K., van der Hoeven Kraft, K. J., & Docktor, J. L. (2021). The Curious Construct of Active Learning. Psychological Science in the Public Interest, 22(1), 8–43. 10.1177/1529100620973974

Luft, J. A., Roehrig, G. H., & Patterson, N. C. (2003). Contrasting landscapes: A comparison of the impact of different induction programs on beginning secondary science teachers’ practices, beliefs, and experiences. Journal of Research in Science Teaching, 40(1), Article 1. 10.1002/tea.10061

Martella, A. M., Lovett, M. C., & Ramsay, L. (2024). Implementing Active Learning: A Critical Examination of Sources of Variation in Active Learning College Science Courses. Journal on Excellence in College Teaching, 32(1), Article 1. https://celt.miamioh.edu/ojs/index.php/JECT/article/view/210

Mazur, E. (1997). Peer instruction: A user’s manual. Prentice Hall.

Mesa, V., & Chang, P. (2010). The language of engagement in two highly interactive undergraduate mathematics classrooms. Linguistics and Education, 21(2), Article 2. 10.1016/j.linged.2010.01.002

Mortimer, E., & Scott, P. (2003). Meaning Making In Secondary Science Classrooms. McGraw-Hill Education (UK).

Neuwirth, E. (2022). RColorBrewer: ColorBrewer Palettes (Version 1.1-3) [Computer software]. https://cran.r-project.org/web/packages/RColorBrewer/index.html

O’Connor, C., & Michaels, S. (2007). When Is Dialogue ‘Dialogic’? Human Development, 50(5), Article 5.

Owens, M. T., Trujillo, G., Seidel, S. B., Harrison, C. D., Farrar, K. M., Benton, H. P., Blair, J. R., Boyer, K. E., Breckler, J. L., Burrus, L. W., Byrd, D. T., Caporale, N., Carpenter, E. J., Chan, Y.-H. M., Chen, J. C., Chen, L., Chen, L. H., Chu, D. S., Cochlan, W. P., … Tanner, K. D. (2018). Collectively improving our teaching: Attempting biology department–wide professional development in scientific teaching. CBE Life Sciences Education, 17(1), ar2.

Pianta, R. C., & Hamre, B. K. (2009). Conceptualization, Measurement, and Improvement of Classroom Processes: Standardized Observation Can Leverage Capacity. Educational Researcher, 38(2), Article 2. 10.3102/0013189X09332374

R Core Team. (2019). *R: A language and environment for statistical computing.* [Computer software]. R Foundation for Statistical Computing.

Reisner, B. A., Pate, C. L., Kinkaid, M. M., Paunovic, D. M., Pratt, J. M., Stewart, J. L., Raker, J. R., Bentley, A. K., Lin, S., & Smith, S. R. (2020). I’ve Been Given COPUS (Classroom Observation Protocol for Undergraduate STEM) Data on My Chemistry Class… Now What? Journal of Chemical Education, 97(4), Article 4. 10.1021/acs.jchemed.9b01066

Rozhenkova, V., Snow, L., Sato, B. K., Lo, S. M., & Buswell, N. T. (2023). Limited or complete? Teaching and learning conceptions and instructional environments fostered by STEM teaching versus research faculty. International Journal of STEM Education, 10(1), Article 1. 10.1186/s40594-023-00440-9

Schunk, D. H. (2012). *Learning Theories: An Educational Perspective*. Pearson.

Sheridan, B. J., & Smith, B. (2020). How Often Does Active Learning Actually Occur? Perception versus Reality. AEA Papers and Proceedings, 110, 304–308. 10.1257/pandp.20201053

Smith, M. K., Jones, F. H. M., Gilbert, S. L., & Wieman, C. E. (2013). The Classroom Observation Protocol for Undergraduate STEM (COPUS): A new instrument to characterize university STEM classroom practices. CBE Life Sciences Education, 12(4), 618–627.

Smith, M. K., Wood, W. B., Krauter, K., & Knight, J. K. (2011). Combining Peer Discussion with Instructor Explanation Increases Student Learning from In-Class Concept Questions. CBE—Life Sciences Education, 10(1), Article 1. 10.1187/cbe.10-08-0101

Sourial, N., Wolfson, C., Zhu, B., Quail, J., Fletcher, J., Karunananthan, S., Bandeen-Roche, K., Béland, F., & Bergman, H. (2010). Correspondence analysis is a useful tool to uncover the relationships among categorical variables. Journal of Clinical Epidemiology, 63(6), Article 6. 10.1016/j.jclinepi.2009.08.008

Stains, M., Harshman, J., Barker, M. K., Chasteen, S. V., Cole, R., DeChenne-Peters, S. E., Eagan, M. K., Esson, J. M., Knight, J. K., Laski, F. A., Levis-Fitzgerald, M., Lee, C. J., Lo, S. M., McDonnell, L. M., McKay, T. A., Michelotti, N., Musgrove, A., Palmer, M. S., Plank, K. M., … Young, A. M. (2018). Anatomy of STEM teaching in North American universities. Science, 359(6383), Article 6383. 10.1126/science.aap8892

Stains, M., & Vickrey, T. (2017). Fidelity of implementation: An overlooked yet critical construct to establish effectiveness of evidence-based instructional practices. CBE Life Sciences Education. 10.1187/cbe.16-03-0113

Streveler, R. A., & Menekse, M. (2017). Taking a Closer Look at Active Learning. Journal of Engineering Education, 106(2), Article 2. 10.1002/jee.20160

Theobald, E. J., Hill, M. J., Tran, E., Agrawal, S., Arroyo, E. N., Behling, S., Chambwe, N., Cintrón, D. L., Cooper, J. D., Dunster, G., Grummer, J. A., Hennessey, K., Hsiao, J., Iranon, N., Jones, L., Jordt, H., Keller, M., Lacey, M. E., Littlefield, C. E., … Freeman, S. (2020). Active learning narrows achievement gaps for underrepresented students in undergraduate science, technology, engineering, and math. Proceedings of the National Academy of Sciences, 117(12), 6476–6483. 10.1073/PNAS.1916903117

Tondeur, J., van Braak, J., Ertmer, P. A., & Ottenbreit-Leftwich, A. (2017). Understanding the relationship between teachers’ pedagogical beliefs and technology use in education: A systematic review of qualitative evidence. Educational Technology Research and Development, 65(3), Article 3. 10.1007/s11423-016-9481-2

Turhan, N. S. (2020). Karl Pearson’s Chi-Square Tests. Educational Research and Reviews, 16(9), Article 9.

Van Booven, C. D. (2015). Revisiting the Authoritative–Dialogic Tension in Inquiry-Based Elementary Science Teacher Questioning. International Journal of Science Education, 37(8), Article 8. 10.1080/09500693.2015.1023868

Vickrey, T., Rosploch, K., Rahmanian, R., Pilarz, M., & Stains, M. (2015). Research-based implementation of peer instruction: A literature review. CBE Life Sci Educ, 14(1), es3.

Wei, T., Simko, V., Levy, M., Xie, Y., Jin, Y., Zemla, J., Freidank, M., Cai, J., & Protivinsky, T. (2021). *corrplot: Visualization of a Correlation Matrix* (Version 0.92) [Computer software]. https://cran.r-project.org/web/packages/corrplot/index.html

Wood, A. K., Galloway, R. K., Sinclair, C., & Hardy, J. (2018). Teacher-student discourse in active learning lectures: Case studies from undergraduate physics. Teaching in Higher Education, 23(7), Article 7. 10.1080/13562517.2017.1421630

Woolley, S. L., Woolley, A. W., & Hosey, M. (1999). *Impact of Student Teaching on Student Teachers’ Beliefs Related to Behaviorist and Constructivist Theories of Learning*. https://eric.ed.gov/?id=ED430964

Zhang, Y. (2009). Classroom Discourse and Student Learning. Asian Social Science, 4(9), Article 9. 10.5539/ass.v4n9p80

