## Supplementary material for "Development of the Follow-Up Discourse Observation Protocol (FUDOP) for characterizing instructor active-learning follow-up behaviors": All supplemental materials

Supplemental Table S1. Dataset details

| | University A | University B | University C |
|---|---|---|---|
| **Public/Private** | Public | Private | Public |
| **Carnegie Classification** | R1 | R1 | R2 |
| **Size** | Large | Medium | Medium |
| **Setting** | Highly residential | Primarily residential | Primarily residential |
| **Enrollment profile** | Majority undergraduate | Majority graduate | Very high undergraduate |
| **Region** | West Coast | Midwest | West Coast |
| **HSI Status** | No | No | Yes |
| **Number of instructors analyzed** | 4 | 8 | 6 |
| **Number of courses analyzed** | 1 | 4 | 2 |
| **Number of course offerings analyzed** | 3 | 8 | 4 |
| **Number of lectures analyzed** | 23 | 23 | 22 |
| **Length of lecture sections** | 50-75 minutes | 50 minutes | 50-75 minutes |

Supplemental Table 2. IRR Percent Agreement

| Code | IRR Percent Agreement |
|---|---|
| Generative | 94.29% |
| Requesting | 88.57% |
| Checking-In | 100.00% |
| Prompting for Individual Thinking | 100.00% |
| Broadening Conversation | 100.00% |
| General Summary | 97.14% |
| Student Answer-Oriented Summary | 97.14% |
| Hinting | 97.14% |
| Announcing Answer | 100.00% |
| Delayed Follow-Up | 100.00% |
| Linking | 97.14% |
| Foreshadowing | 94.29% |
| Soliciting Individual Student's Response | 97.14% |
| Moving On | 100.00% |

Supplemental Table #3: Original FUDOP Coding Scheme and Expert Panel Feedback. Follow-up is what the instructor does after an active learning activity has occurred in class in response to that activity. Multiple codes can be used to code for one follow-up. A follow-up segment ends when the instructor switches to a new slide to discuss another topic or initiates a new active learning activity.

| **Code** | **Source** | **Original Description** | **Original Example Dialogues** | **Expert Panel Feedback** | **New Description (Edit is in Blue)** | **New Example Dialogues (Edit is in Blue)** |
|---|---|---|---|---|---|---|
| **Student-centered** | Whole class participates | | | | | |
| **Generative** | **CDOP** | Prompting **close-ended** questions to the whole class (requires student input); no explicit explanation given. | Instructor: "Do we see a dark smudge on this gel?" Whole class: "Yes." Instructor: "How many DNA bands?" Whole class: "Two." | Clarify "no explanation/ requires student input". | Prompting **close-ended** questions (can be answered with a brief response, often a simple "yes" or "no" or a specific piece of information) to the whole class (requires student input); no explicit explanation given. | Instructor: "Do we see a dark smudge on this gel?" Whole class: "Yes." Instructor: "How many DNA bands?" Whole class: "Two." |
| **Requesting** | **CDOP** | Asking **open-ended** questions to the whole class (not soliciting an individual volunteer, does not require an answer from the class). | Instructor: "What does everybody think?" | Say to the entire class; specify that it does not need to be answered. Maybe change the example dialogue to explain student response. Is there also an example that is content-based or content-focused like above? | Asking the whole class **open-ended** questions (e.g. prompting a more elaborate and detailed response, encouraging the student to express their thoughts and opinions) (not soliciting an individual volunteer). | Instructor: "What are some possible reasons why we have a decrease in translation of Protein X? What does everybody think?" |
| **Checking-In** | **CDOP** | Checking in with students to **make sure they are understanding** what's happening or see how they are feeling. | Instructor: "Does that make sense, any Q's?" | In COPUS one of the most confusing thing is whether this kind of question is rhetorical or not. When it is rhetorical, it does not count in COPUS, although deciding if something is rhetorical or not is kind of hard. | Checking in with students to **make sure they are understanding** what's happening or see how they are feeling. | Instructor: "Does that make sense, any Q's?" |
| **Prompting for Individual Thinking** | New | Giving students the opportunity to **think about questions/concepts individually.** | Instructor: "Think about this question individually." | This and next one seem "physical" compared to the above which are more cognition focused, perhaps change wording. | Giving students the opportunity to **think about questions/concepts individually.** | Instructor: "Think about this question individually." |

| | | | | | | |
|---|---|---|---|---|---|---|
| **Broadening Conversation** | **CDOP "Explaining"** | Telling them to **talk among their neighbors** (must change behavior; cannot be a reminder). | Instructor: "Discuss with your neighbors now." | "Student" must change behavior; this was confusing to some panelists. | Asking students to **talk among their peers** (students must change behavior; cannot be a reminder if the activity itself was already peer discussion). | Instructor: "Discuss with your neighbors now." |
| **Instructor-centered** | One or no student participates | | | | | |
| **Student Answer-Oriented Summary** | **CDOP "Evaluating"** | **Giving summary based on student responses** (ie."most people picked XYZ" and explains it based on XYZ). | Instructor: "Our class is divided between B and C." Instructor explains the question based on choices B and C (student response). | Specify that it is an explanation specific to the students' answers– maybe change "summary" to explanation"; reword "class is divided". | **Giving explanations based on student responses** (ie."most people picked XYZ" and explains it based on XYZ). | Instructor: "Our class is evenly split between choices B and C." Instructor explains the question based on choices B and C (student response). |
| **General Summary** | **CDOP "Sharing"** | **Summarizing the answer**; explanation of question w/o explicitly referring to student responses. | Instructor: "The correct answer is X because…" | | **Explaining the answer**; explanation of question w/o explicitly referring to student responses. | Instructor: "The correct answer is X because…" |
| **Hinting** | **CDOP "Sharing"** | **Providing guidance**; not giving out answers. | Instructor: "Consider point X when trying to answer this question…" | | **Providing guidance on how to answer the question** without giving out answers. | Instructor: "Consider point X when trying to answer this question…" |
| **Announcing Answer** | New | Only **telling students the answer,** without explanation. | Instructor: "The correct answer is C. Our next topic is…" | | Only **telling students the answer,** without explanation. | Instructor: "The correct answer is C. Our next topic is…" |
| **Delayed Follow-Up** | New | **Following up throughout lecture** or in the next lecture. | Instructor: "I will come back to this question later." | Foreshadowing vs delayed follow-up, clarify the time difference? | **Not following up immediately; instead following up later in lecture** or in upcoming lecture. | Instructor: "I will hold off discussing this question until next class." |
| **Linking** | **CDOP** | **Associating current topic to past topic or previous mentions, recalling information** (explicit only). | Instructor: "As we talked about last week/yesterday…" | Linking vs recalling, explicit definition, may not know how different a concept is from another. | **Associating current topic to past topic or previous mentions, recalling information** (explicit only). | Instructor: "As we talked about last week/yesterday…" |

| | | | | | | |
|---|---|---|---|---|---|---|
| **Fore-shadowing** | **CDOP** | **Associating current topic to future topic** (explicit only). | Instructor: “We will be discussing … next week/ upcoming lecture” | Foreshadowing vs delayed follow-up: time difference? | During follow-up, **associating current topic to future topic** (explicit only). | Instructor: “We will come back to this idea again when we discuss DNA transcription next week.” |
| **Soliciting Individual Student's Response** | New | **Asking for individual student(s) to answer the question** by raising their hand. | Instructor: “Can we have a volunteer answer this?”<br>One student answers | | **Explicitly asking for individual student(s) to answer the question** while addressing the whole class. | Instructor: “Can we have a volunteer answer this?”<br>One student answers. |
| **Other** | | | | | | |
| **Moving On** | **CDOP** | Doing nothing, **just moving on** (no follow up). | Instructor: “Let’s move on to the next topic” (and then moves on to next question/topic) | More specific example so it doesn’t look like announce answer; definition could state that no explanation is given. | Doing nothing, **just moving on** (no follow up). | Instructor: “Let’s move on to the next topic” (and then moves on to next question/topic) |

Supplemental Table 4. Between-Instructor p-values (continue from Figure 4)

University A

| Code | P-Value | |
|---|---|---|
| Generative | 0.0186 | |
| Requesting | 0.0006 | * |
| Checking-In | 0.0000 | * |
| Prompting for Individual Thinking | N/A | |
| Broadening Conversation | 0.4853 | |
| General Summary | 0.0402 | |
| Student Answer-Oriented Summary | 0.0181 | |
| Hinting | 0.0169 | |
| Announcing Answer | 0.0000 | * |
| Delayed Follow-Up | 0.3162 | |
| Linking | 0.2851 | |
| Foreshadowing | 0.4252 | |
| Soliciting Individual Student's Response | 0.6118 | |
| Moving On | 0.9718 | |

University B

| Code | P-Value | |
|---|---|---|
| Generative | 0.3682 | |
| Requesting | 0.6731 | |
| Checking-In | 0.5398 | |
| Prompting for Individual Thinking | N/A | |
| Broadening Conversation | 0.4866 | |
| General Summary | 0.4483 | |
| Student Answer-Oriented Summary | 0.0005 | * |
| Hinting | 0.7340 | |
| Announcing Answer | 0.3151 | |
| Delayed Follow-Up | NaN | |
| Linking | 0.5330 | |
| Foreshadowing | 0.6260 | |
| Soliciting Individual Student's Response | 0.4292 | |
| Moving On | N/A | |

University C

| Code | P-Value | |
|---|---|---|
| Generative | 0.2811 | |
| Requesting | 0.4192 | |
| Checking-In | 0.1637 | |
| Prompting for Individual Thinking | 0.7719 | |
| Broadening Conversation | 0.5089 | |
| General Summary | 0.0130 | |
| Student Answer-Oriented Summary | 0.0013 | * |
| Hinting | 0.0936 | |
| Announcing Answer | 0.6316 | |
| Delayed Follow-Up | 0.8878 | |
| Linking | 0.8898 | |
| Foreshadowing | 0.8793 | |
| Soliciting Individual Student's Response | 0.0176 | |
| Moving On | N/A | |

Supplemental Figure 1. Sample coding.

### FUDOP coding example #1

0:02
Instructor: The class was split 50-50. Can those who answered B give me a reason? It's good that so many people answered B. Why did you pick B? Oh yeah?

0:46
[An individual student raises their hand, the instructor calls on them, and the individual student gives an answer.]

Soliciting Individual Student's Response

1:08
Instructor: So let me see if I get your reason. You think the enhancer may or may not be needed for transcription. Is that correct? How about we put it in a different way? What does the 'enhancer' mean- to enhance something?

1:35
Instructor: So if we are talking about transcription, the only meaning for "enhancer" is to enhance transcription. So enhancer A is a DNA sequence, DNA element, usually 68 to 80 nucleotides long. Very Short. They are there to enhance transcription.

1:59
Instructor: How can a piece of DNA that enhances transcription occur at the promoter? That's your question, right? You think well, because the promoter is required for transcription, enhancement may or may not be required for transcription. That is correct.

Linking

2:23
Instructor: Remember the CAP protein we talked about the last time- you got CAP binding to the binding site, the promoter is downstream, then you push it on the polymerase to activate transcription. So you need to bind to something in order to activate transcription, right? An activator. So, an activator is a transcription factor, a positive regulator of transcription- that means it activates transcription. It activates transcription.

2:55
Instructor: If you don't have an enhancer or the enhancer cannot bind to activator, what will happen to transcription?

3:03
Class choral response: Decrease.

Generative

3:06
Instructor: Does that make sense? Good?

Checking-in

| 1. Student Centered | | | | | 2. Instructor Centered | | | | | | | | 3.Others |
|---|---|---|---|---|---|---|---|---|---|---|---|---|---|
| Generative | Requesting | Checking-In | Prompting for Individual Thinking | Broadening Conversation | General Summary | Student Answer-Oriented Summary | Hinting | Announcing Answer | Delayed Follow Up | Linking | Foreshadowing | Soliciting Individual Student's Response | Moving On |
| | | 1 | | | | 1 | | | | 1 | | 1 | |

### FUDOP coding example #2

18:17
Instructor: Well, the answers are all over the place.

Broadening Conversation

18:22
Instructor: So I want you guys to, to discuss this and then we're gonna be, it seems that you can, you can the question.

18:35
Instructor: So let's take a look at this.

18:38
Instructor: So we have two choices, either be identified by conation or we identified by and this packaging is either dominant or excessive for again, I OK.

19:17
Instructor: I'm gonna close the poll in 10 seconds.

19:32
Instructor: So the number of people chose one and the number of people chose C

Student Answer-Oriented Summary

19:39
Instructor: So, or a right, we, we didn't use any hybridization in this, in this. We're not, you don't know what the team is. The only way we can identify this gene is by function, in this case, it's a gain of function that is essentially complementation. so, we're not restoring the wild-type phenotype. In this case, we're providing these cells with a new function. But nonetheless, this is essentially complementation, but it's, it's a data function and the function that we are providing them with is the ability to grow uncontrolled. The second thing is that this be patient, dominant or recession. So our wild type fiberglass that the wild type genotype, the wild type genes were dominant, then we would never get this uncontrolled growth deal. And so our accomplish must be done. So it's providing a new function for these cells, the ability to grow in an uncontrolled fashion. The right answer is a we essentially close this team by complementation, but it, it's a little bit different than we normally do. When we do complementation, we restore the wild type phenotype. In this case, what we're doing is we're giving these cells, a new phenotype, the ability to grow in an uncontrolled fashion. And the only way that we would be able to identify it is if this gene is dominant because it has to provide this new function if you like in the face of the wild type genes that are already present.

(continue)

21:35
Instructor: So any questions about the logic here, go ahead.

Checking-in

21:41
Instructor: So the complementation is when we identify something by virtue of its, by virtue of its activity.

21:50
Instructor: So in this case, can we confer uncontrolled growth on itself?

21:56
Instructor: So we do hybridization to a library. In that case, we must already have the seats. So that would be a case that I identified the gene as per I'm interested in seeing whether the same gene exists in a mouse library. I generate a probe that corresponds to the fly gene and I use that to identify the mouse. I have no idea what the is I have to identify by his ability to confer a function on the self that I'm studying.

22:30
Instructor: Any other questions?

22:33
Instructor: All right.

| 1. Student Centered | | | | | 2. Instructor Centered | | | | | | | | 3.Others |
|---|---|---|---|---|---|---|---|---|---|---|---|---|---|
| Generative | Requesting | Checking-In | Prompting for Individual Thinking | Broadening Conversation | General Summary | Student Answer-Oriented Summary | Hinting | Announcing Answer | Delayed Follow Up | Linking | Foreshadowing | Soliciting Individual Student's Response | Moving On |
| | | 1 | | 1 | | 1 | | | | | | | |

Supplemental Figure 2. Scree plot for Figure 3A.

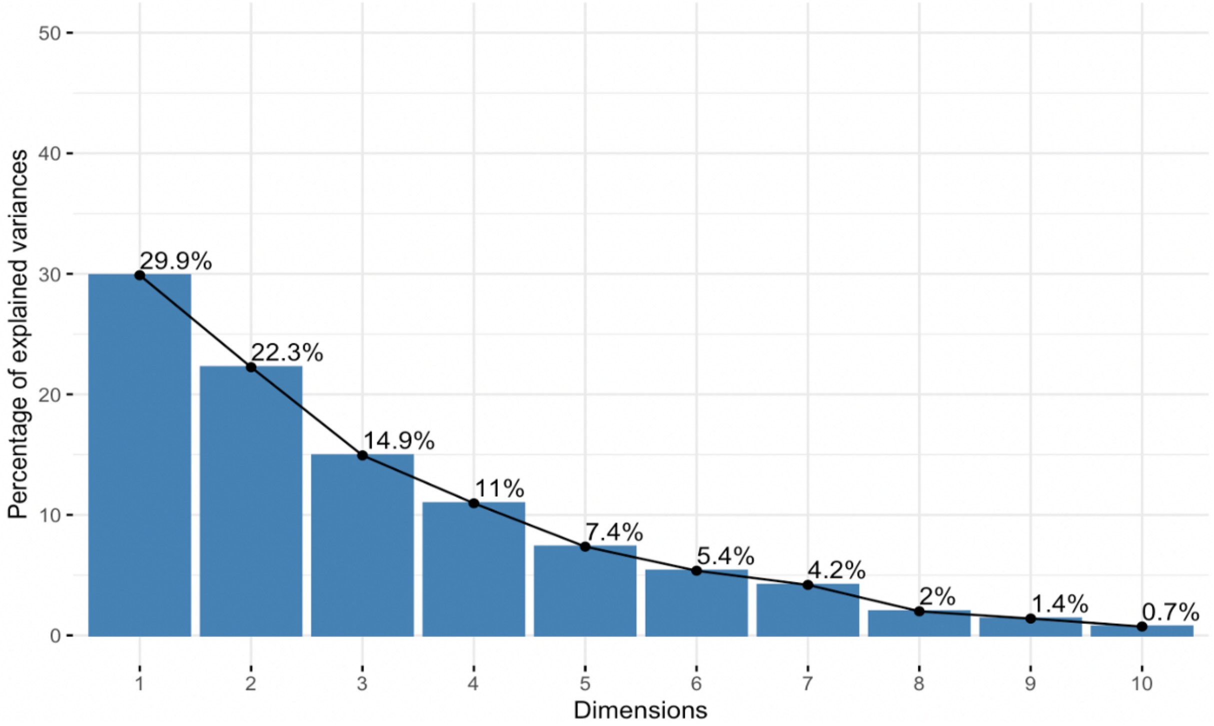

Supplemental Figure 3. Variability of FUDOP Codes After Single Active Learning Segment.

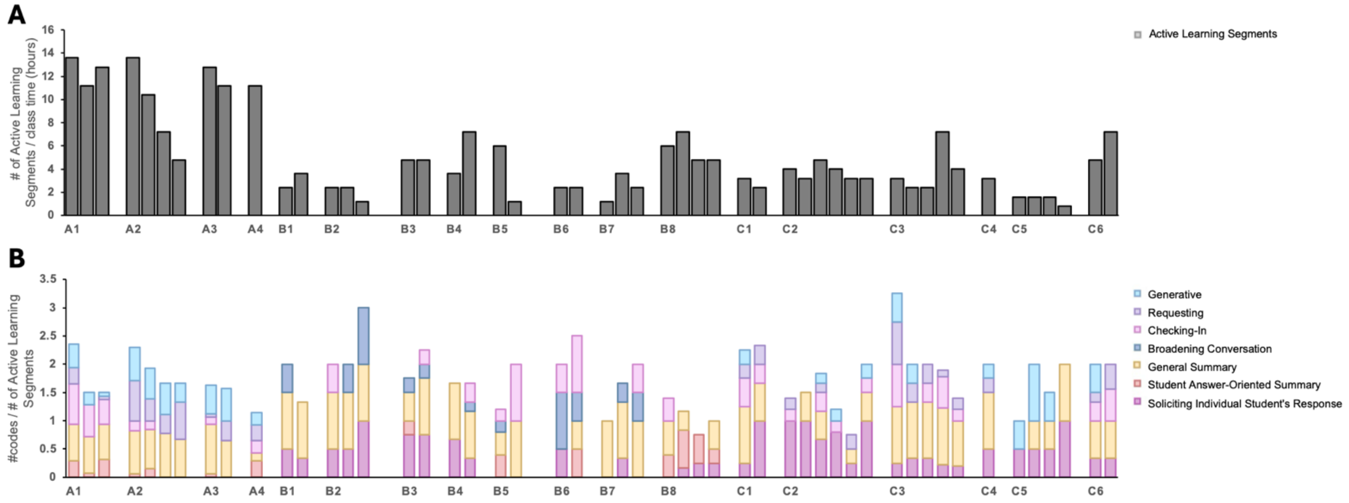
